# Evaluation of Prostaglandin Receptor Agonists and Eupatilin in the Context of Nephronophthisis

**DOI:** 10.1101/2025.01.15.633139

**Authors:** Alice Tata, Guillaume Rocha, Marguerite Hureaux, Alice Serafin, Esther Porée, Lucie Menguy, Nicolas Goudin, Nicolas Cagnard, Lilian Gréau, Marc Fila, Luis Briseno-Roa, Jean-Philippe Annereau, Sophie Saunier, Alexandre Benmerah

## Abstract

**Background:** Primary cilia are sensory antennas that are present on the majority of quiescent vertebrate cells where they mediate key signaling during development and in response to environmental stimuli. Defects in primary cilia result in a group of heterogeneous inherited disorders with overlapping phenotypes, called ciliopathies. Nephronophthisis is an autosomal recessive tubulo-interstitial kidney ciliopathy with more than 25 identified genes called *NPHP*. Presently, no treatment exists beyond supportive care and kidney transplant, underscoring the need for novel therapies.

**Methods:** Using a phenotypic screening approach in cultured cell lines, we previously identified prostaglandin analogues as candidate therapeutic molecules based on their ability to rescue ciliogenesis defects in kidney tubular cells from *NPHP1* patients. Here, we have investigated the potential beneficial effects of ROCK inhibitors and Eupatilin, similarly identified by other groups in different NPHP contexts, in kidney cells from *NPHP1* and *IQCB1/NPHP5* patients as well as in a zebrafish *NPHP* mutant line (*traf3ip1/ift54*).

**Results:** Eupatilin partially rescued *NPHP1*-associated ciliogenesis defects. Transcriptomic analyses pointed out that cell cycle progression was inhibited by Eupatilin, likely explaining its broad effects on cilia assembly. Interestingly, while ciliary defects also observed in *NPHP5* patient cells were rescued by both prostaglandins and Eupatilin, only prostaglandin analogues were able to reduce pronephric cysts size in the used *nphp* zebrafish model.

**Conclusion:** Our study indicates that these molecules can show beneficial effects across genetic contexts and shed light on their potential as therapeutic interventions for nephronophthisis.

## Introduction

Nephronophthisis (NPH; OMIM: #256100) is a tubulo-interstitial hereditary kidney disease caused by variants in genes encoding proteins functioning at different compartments of the primary cilium (PC), a cellular signaling hub found on most vertebrate cells including kidney epithelial cells. It is therefore classified among ciliopathies, a complex group of disorders caused by PC dysfunction affecting various organs including kidney, retina, central nervous system and skeleton^1–3^.

PC are assembled in quiescent cells (G_0_) from the centrosome. During this complex process, the mother centriole docks onto the plasma membrane, then called the basal body, and elongates to form the microtubule-based axoneme which is wrapped into the ciliary membrane. The specific protein and lipid composition of the cilium is maintained by the transition zone (TZ), a structure at the base of PC which prevents free exchange of proteins between the cytoplasm and the ciliary compartment. Most ciliary components, including ciliary membrane proteins, must therefore be actively transported by the IFT (intraflagellar transport) which selects cargos and transport them in and out along the axoneme. This process is required for ciliogenesis as well as for PC-dependent signaling which is based on the ciliary localization of specific sets of receptors and their downstream effectors^1^.

To date, more than 25 genes have been linked to NPH (*NPHPs*) which encode proteins that are playing important roles in PC-mediated signaling functions and/or ciliogenesis^1–3^. The most frequent one is *NPHP1* accounting for nearly 25% of NPH cases, mostly harboring a homozygous deletion of the gene. NPHP1 plays an important role at the TZ together with many other *NPHP* gene products^3^. Three main clinical subtypes have been described for NPH. While they all lead to end stage kidney disease (ESKD), they differ by the age of onset as well as by the prevalence of cysts. Juvenile NPH with a mean age of onset of 13 years is by far the most frequent form, accounting for half of NPH cases and for 5 to 10% of ESKDs in children and young adults. Biallelic loss of function (LOF) or missense variants in *NPHP1*, *NPHP4* and *NPHP5/IQCB1* are among the main cause juvenile/late onset NPH which can be isolated (*NPHP1*, *NPHP4*) or associated with retinal dystrophies in Senior-Løken syndrome (*NPHP1*, *NPHP4*, *NPHP5;* OMIM: #266900)^3,4^.

There is no treatment available for NPH which inevitably leads to ESKD requiring dialysis and kidney transplant. Molecules aiming at limiting cyst growth were discovered based on studies made in cystic infantile NPH mouse models. However, those molecules are unlikely to be relevant in juvenile NPH since kidney cysts are mainly observed in the latest stages of the disease^5,6^. Interestingly, NPH-causing variants often lead *in vitro*, depending on the cell type, to a decreased proportion of ciliated cells. As ciliogenesis can be easily quantified by immunofluorescence image analyses, we and other groups used this rationale to identify compounds able to rescue the ciliogenesis defects observed in the context of NPH^5,6^. Using this approach, we recently identified prostaglandin E1 (PGE1) and other agonists of PGE2 receptors (EPs) as a class of molecules that restores ciliary defects *in vitro* as well as kidney and retina-associated phenotypes *in vivo* in the context of *NPHP1*^7^. Other groups have similarly identified additional candidate therapeutic molecules including ROCK inhibitors and Eupatilin, in the context of *NPHP8* and *NPHP6*, respectively^5,6,8^. The aims of the present study were to further characterize hits that we identified in our initial screen and to evaluate whether if compounds identified in a given *NPHP* context could show broader efficiency in different genetic backgrounds. To achieve these aims, the effects of those compounds were tested in kidney tubular cells from *NPHP1* and *NPHP5* patients as well as in a zebrafish model of NPH.

## Results

### Eupatilin and ROCK inhibitor increase Cilia incidence in NPHP1 patient URECs

A phenotypic screening approach previously led to the identification of small molecules able to modulate the ciliopathy-associated phenotypes. Among them, Alprostadil/PGE1, a prostaglandin analogue, robustly rescued ciliogenesis in *NPHP1* patient kidney epithelial cells^7^. Besides prostaglandins, other hits from this screen remained to be fully investigated including statins, antibiotics (cephalosporins), glucocorticoids as well as other compounds known to modulated cAMP signaling (Table 1). Those molecules were selected based on their published or potential effects on ciliogenesis. In addition, Eupatilin and ROCK inhibitors were also included based on their described positive effects on the ciliogenesis defects in the context of NPH^5,6,8^ and the observed increased RHO activity in *NPHP1* patient cells^7^. Those molecules were further tested on immortalized urinary-derived renal epithelial cells (URECs) which were obtained from control and NPH individuals presenting with homologous deletion of *NPHP1*^7^.

**Table 1.**

| Compounds | Class | Known/*expected* effect on cilia |
| --- | --- | --- |
| Alprostadil | EP agonist | Rescue cilionogenesis in NPHP1 URECs<sup>7</sup> |
| Taprenepag | EP2 agonist | *Rescue ciliogenesis in NPHP1 URECs* |
| Y-27632 | Rock inhibitor | Positive effect on cilia in ciliopathy conditions<sup>5,18</sup> |
| Dexamethasone | Glucocorticoid | Positive effect on cilia<sup>41–43</sup> |
| Isoproterenol | β-adrenergic agonist | *Positive effect on cilia through cAMP?* |
| Theophylline | Phosphodiesterase inhibitor | *Positive effect on cilia through cAMP?* |
| Fluvastatin | HMG-CoA reductase inhibitor | Positive effect on cilia in high cholesterol conditions<sup>39,40</sup> |
| Simvastatin | HMG-CoA reductase inhibitor | Positive effect on cilia in high cholesterol conditions<sup>39,40</sup> |
| Cefotetan | Cephalosporin | Positive effect on cilia in cancer cells<sup>41</sup> |
| Cefaclor | Cephalosporin | Positive effect on cilia in cancer cells<sup>41</sup> |
| Eupatilin | Flavonoid | Positive effect on cilia in ciliopathy conditions<sup>16,17,50</sup> |

Ciliation was quantified using the Harmony software (see methods) based on ARL13B (ciliary membrane) and γ-Tubulin (basal body) stainings (Figure 1A). As previously shown^7^, *NPHP1* URECs displayed reduced PC incidence compared to control (Figure 1A,B) with shorter cilia (Figure 1C). The same *NPHP1* cell lines were treated for 48 hours with each of the selected hit compounds using Alprostadil (Alpro) as a positive control. Unfortunately, none of the hits coming from the screen showed a positive effect on ciliation (Supplementary Figure S1). Interestingly, treatment of *NPHP1* URECs with Eupatilin increased ciliation (Figure 1D-F) similarly as previous findings in *NPHP6* conditions^16,17^. In addition, treatment with the ROCK inhibitor Y-27632 was also able to increase ciliation in *NPHP1* URECs (Figure 1E,G). Notably, ciliation was not increased in control cells treated with those two compounds (Figure 1F,G).

**Figure 1:**
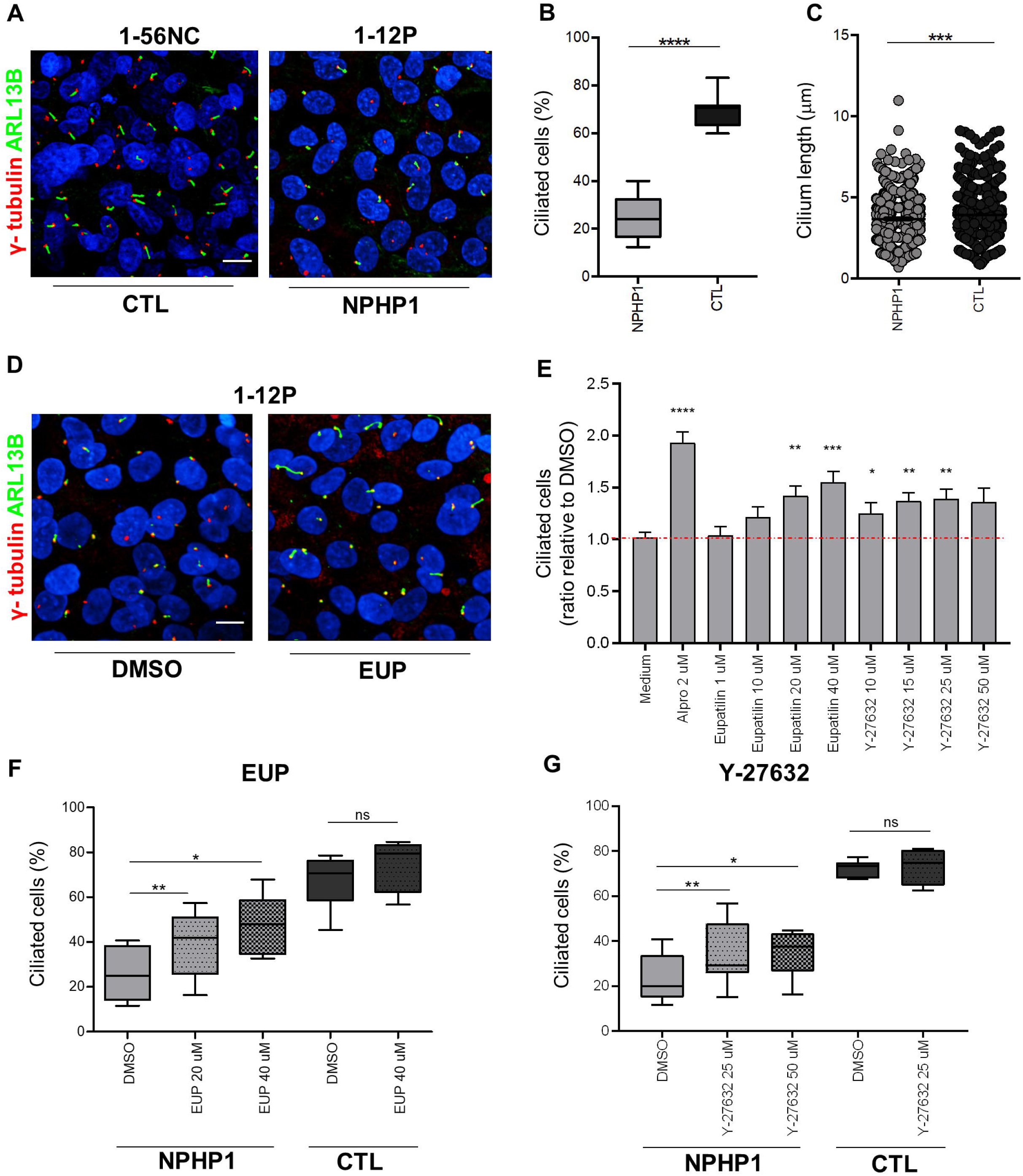
Eupatilin and ROCK inhibitors rescue ciliogenesis defects in *NPHP1* URECs. **(A)** Control (1-56NC) and *NPHP1* (1-12P) URECs grown in ciliogenesis conditions for 5 days at non permissive temperature (39°C) were fixed and stained for primary cilia (ARL13B, green) and basal bodies (γ-tubulin, red)**. (B)** Ciliogenesis in *NPHP1* (1-12P and 1-03P) and CTL (1-56NC) URECs was quantified using Harmony software and expressed as percentage of ciliated cells shown using a box-and-whisker plot (n=5 to 11 experiments). Mixed linear-regression model with quasibinomial penalization taking into account the correlation of observations coming from the same individuals using a random effect on the cell line: ****P<0.0001. **(C)** Cilium length in *NPHP1* (1-12P and 1-03P) and CTL (1-56NC) URECs length was quantified as explained in Methods and expressed in μm using a dot-plot (n=3 experiments). Unpaired two-tailed Student’s t-test: ***P<0.001. **(D)** *NPHP1* URECs (1-12P) treated with either DMSO (0,04%) or 20μM EUP for 48h were fixed and stained as in (**A**) for primary cilia (ARL13B, green) and basal body (γ-tubulin, red). **(E)** Ciliogenesis in 1.12P and 1.03P *NPHP1* URECs treated for 48h with either Alprostadil (Alpro), Eupatilin (EUP) or ROCK inhibitor (Y-27632) was expressed as the ratio to control DMSO-treated cells. Mean ± SEM (n=4 to 7 experiments); paired two-tailed Student’s t-test: ****P<0.0001, ***P<0.001, **P<0.01, *P<0.05 **(F, G)** Ciliogenesis in URECs derived from CTL (1-56NC) and *NPHP1* patients (1-12P and 1-03P) treated for 48h with DMSO or EUP (20 and 40μM; n=4 to 6 experiments per cell line; **F**) or Y-27632 (25 and 50 μM; n= 2 to 5 experiments per cell line) was quantified similarly as in **B**. *P<0.05,**P<0.01, ns: not significant. Scale bars 10μm.

Altogether, those results did not confirm the beneficial effects on ciliogenesis of the compounds identified in our screen on cell lines, they however show that both Eupatilin and ROCK inhibitor can rescue ciliation defects observed in *NPHP1* patient cells.

### Eupatilin modulates the expression of cell cycle and membrane trafficking pathways

While the effects of ROCK inhibitors in the context of PC and renal ciliopathies is well documented^5,18^ the targets and downstream effects of Eupatilin remain poorly understood. Therefore, bulk RNA sequencing (RNAseq) was then performed on the URECs treated with either DMSO or 20μM Eupatilin for 24 hours in ciliogenesis conditions.

Clustering of the samples by significant differential expression (PCA analysis) showed that the Eupatilin-treated samples had similar transcriptome profiles, distinct from DMSO-treated ones (Figure 2A); 4585 genes were differentially expressed (false discovery rate (FDR)-adjusted p-value cutoff of 0.05), with 2041 being downregulated and 2544 upregulated (Supplementary File S1). Metascape was used to analyze the pathways modulated in response to Eupatilin (Figure 2B). Based on the adjusted p-value, the most significantly downregulated gene sets were related to cell-cycle progression (cell cycle, DNA metabolic process, cell cycle checkpoints). In the G_1_-S subset (Supplementary Figure S2A and File S2), there was a decreased expression of cyclin-dependent kinase (*CDK2*, *CDK4-6*) as well as of promoters of cell-cycle progression to S phase (*CDC25A, MYBL2* and *RB1*). In the G_2_-M subset (supplementary Figure S2B and File S2), markers of chromosome alignment (*NUP188*, *KIF22*), cilium resorption (*KIF24*, *NEK2*), as well as genes related to chromosome organization and DNA repair (*PCNA*, *ILF2*) were downregulated. Interestingly, Eupatilin upregulates the expression positive regulators of quiescence and/or genes known to be expressed in G_0_-G_1_ cells (*CDKN2B*, *SERINC* and *LARP1*; Supplementary Figure S2C and File S2). RNAseq results were mostly confirmed by qRT-PCR analysis (Figure 2C,D) indicating that Eupatilin treatment likely results in anti-proliferative effects as previously reported^19–21^.

**Figure 2:**
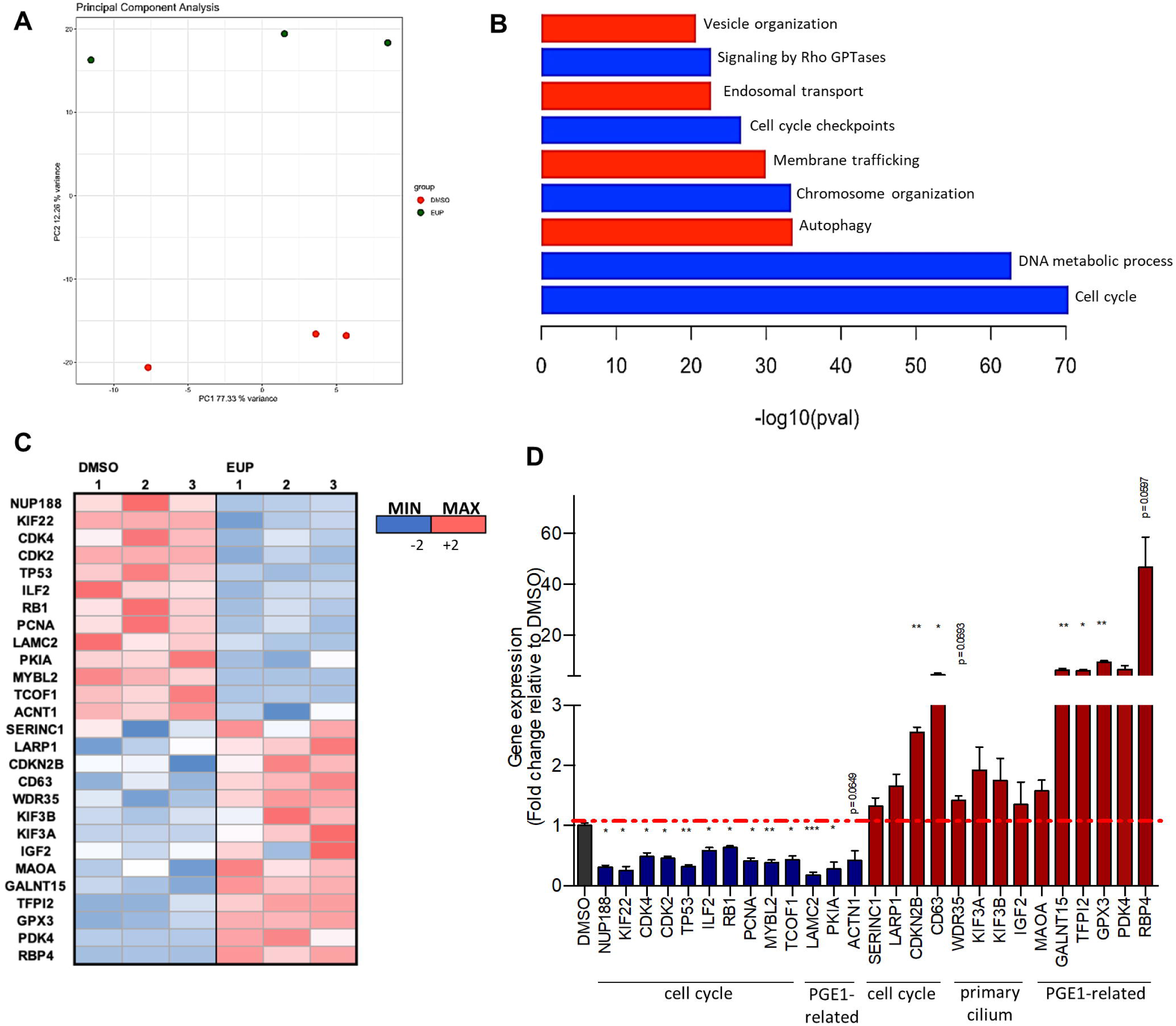
Transcriptomic analyses reveal the cellular pathways modulated in response to Eupatilin treatment. **(A)** PCA analysis showing separation between the EUP treated URECs and the DMSO treated by PC2 variance. **(B)** Pertinent down (blue) or up (red) regulated pathways or relevant processes involving ciliary modulators dysregulated in *NPHP1* URECs upon 24h 20μM EUP treatment were highlighted using Metascape. **(C)** Heatmap generated from qPCR results of 27 selected genes from RNA-seq dataset of *NPHP1* URECs DMSO versus EUP. **(D)** RT-qPCR validating the effect of EUP treatment on the expression of 27 selected dysregulated genes in *NPHP1* URECs. Mean ± SEM (n=3 experiments). Paired two-tailed Student’s t-test:***P<0.001, **P<0.01, *P<0.05, ns: not significant. Bars indicate mean ± SEM.

In addition to cell cycle progression, Eupatilin treatment also downregulated RHO GTPases signaling as well as actin-cytoskeletal protein encoding genes (Figure 2B; Supplementary Figure S3A and File S2), in agreement with its positive effect on ciliation. Interestingly, we previously show that RHO activity was downregulated in *NPHP1* URECs upon Alprostadil/PGE1 treatment^7^. Concerning the upregulated gene families, the most statistically significant enlightened by Metascape was “vesicular trafficking related to autophagy” (Figure 2B; Supplementary Figure S3B and File S3) which was linked to the early steps of ciliogenesis^22^.

In addition to this unbiased pathway analysis, we also focused on specific gene families of interest including ciliary genes and genes that we found modulated upon prostaglandin treatment^7^. A list of ciliary genes was established based on Ciliacarta^23^. Among known positive modulators of ciliogenesis, key IFT components including *IFT88*, *WDR19*, *WDR35*, *WDR11*, *KIF3A* and *KIF3B*, were upregulated (Supplementary Figure S4 and File S4) but this increased expression could not be confirmed by qRT-PCR (Figure 2C,D). Finally, among the genes that were found to be modulated by Alprostadil^7^, 169 were similarly affected by Eupatilin (Supplementary Figure S5A and B and File S4), with some downregulated (*LAMC2*, *PKIA*, *CAV1*, *ACTN1*) and others upregulated (*PDK4*, *ATOH8*, *CDKN2B*), some of which were confirmed by qRT-PCR (Figure 2C,D), indicating a partially shared signature with Alprostadil.

Altogether, our RNAseq analysis confirmed previous studies indicating the cytostatic effects of Eupatilin which therefore may increase ciliogenesis through cell-cycle regulation favoring entry/persistence of cells in G_0_.

### NPHP5 URECs presents ciliogenesis defects similar to NPHP1 which are partially rescued by Alprostadil/PGE1 and Eupatilin

In order to evaluate the potential broader effects of the molecules identified in the context of *NPHP1,* we next focused on *IQCB1/NPHP5* which was previously associated with Senior-Løken syndrome with juvenile NPH^24–29^. URECs were collected from two non-twin brothers (2-05P1 and 2-05P2) harboring compound heterozygous variants in *NPHP5*^29^. Sanger sequencing confirmed the presence of the two LOF variants in URECs which result in decreased *NPHP5* expression (Figure S6).

Ciliogenesis was then quantified in the two *NPHP5* UREC lines. As expected from previous works^30,31^, *NPHP5* URECs from both patients presented severe ciliogenesis defect and shorter cilia (Figure 3A-C). Besides ciliogenesis, *NPHP5* URECs also showed partial TZ defects with decreased staining for TMEM67/NPHP11 (Figure 3D and E) while NPHP4 remained similar as in controls (Supplementary Figure S7A,B). As a potential consequence of the observed TZ defects, the ciliary fluorescence intensity for both ADCY3 and INPP5E, two widely used ciliary membrane markers, was decreased in *NPHP5* URECs compared to control (Figure 3F,G; Supplementary Figure S7C-E).

**Figure 3:**
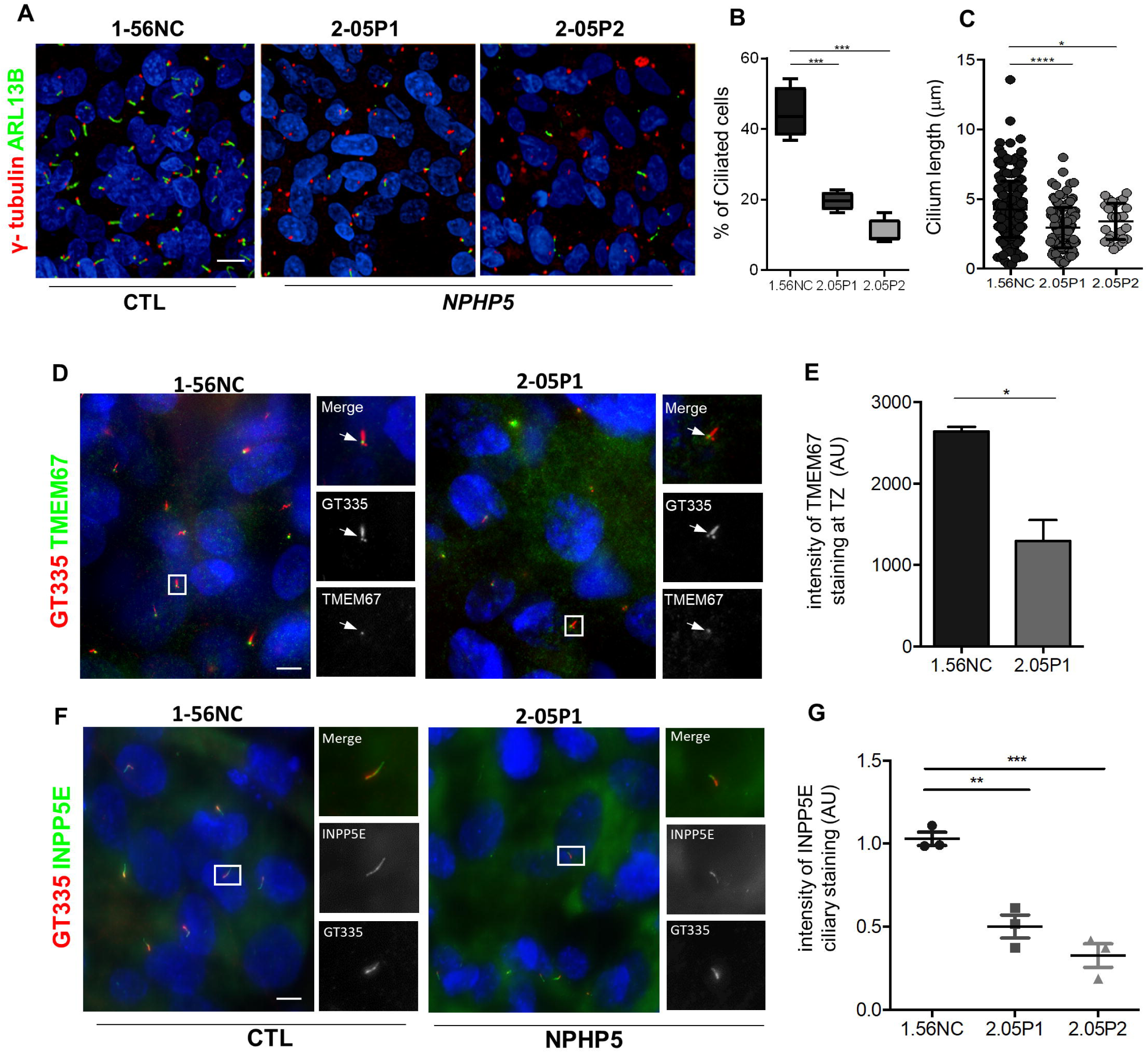
*NPHP5* URECs show ciliogenesis and ciliary composition defects. **(A-C)** Control (1-56NC) and *NPHP5* URECs (2.05P1, 2.05P2) grown in ciliogenesis conditions fixed and stained for primary cilia (ARL13B, green) and basal bodies (γ-tubulin, red) markers (**A**). Ciliogenesis (**B**) and cilium length (**C**) were quantified as in Figure1. **(B**) n=4 experiments; mixed linear-regression model with quasibinomial penalization: ***P<0.001. (**C**) n=3 experiments; unpaired two-tailed Student’s t-test: ****P<0.0001; *P<0.05. **(D)** Ciliated control (1.56.NC) and *NPHP5* (2.05P1) URECs were fixed and stained for basal body and axoneme (GT335, red) and for TMEM67 (green) which is a marker of the transition zone (arrows). **(E)** TMEM67 staining intensity at TZ was quantified as detailed in Methods. n=2 experiments; unpaired two-tailed Student’s t test: *P<0.05. **(F)** Ciliated control (1.56.NC) and *NPHP5* (2.05P1) URECS were fixed and stained for basal body and axoneme (GT335, red) and for INPP5E (green). **(G)** Intensity of INPP5E staining in cilia was quantified as explained in Methods. n=3 experiments; unpaired two-tailed Student’s t test: \*\**P* < 0.01, \*\*\**P* < 0.001. Scale bars 10μm.

In conclusion, URECs from two related patients with LOF variants in *NPHP5* presented with ciliogenesis and ciliary composition defects similar to results obtained in *NPHP1* conditions. We therefore tested prostaglandin receptor agonists, Eupatilin and ROCK inhibitor. In addition to Alprostadil which is able to activate all EPs, Taprenepag (Tap), a selective agonist of EP2^32^ which is highly expressed in URECs, has proved its effects in rescuing the retinopathy phenotype in *Nphp1* knockout mice^7^. In *NPHP5* URECs, both agonists showed positive effects on ciliogenesis, with a similar efficiency as the one observed in *NPHP1* cells (Figure 4A,B), and similarly increased ciliary length (Figure 4C,D). Finally, we also tested the two other compounds which showed a positive effect on ciliogenesis in *NPHP1* URECs (Figure 1). While the ROCK inhibitor Y-27632 did not show significant effect (Figure 5A,B), Eupatilin significantly increased ciliogenesis in *NPHP5* URECs but only at 40μM (Figure 5C,D; see Figure 1F for comparison with *NPHP1*).

**Figure 4:**
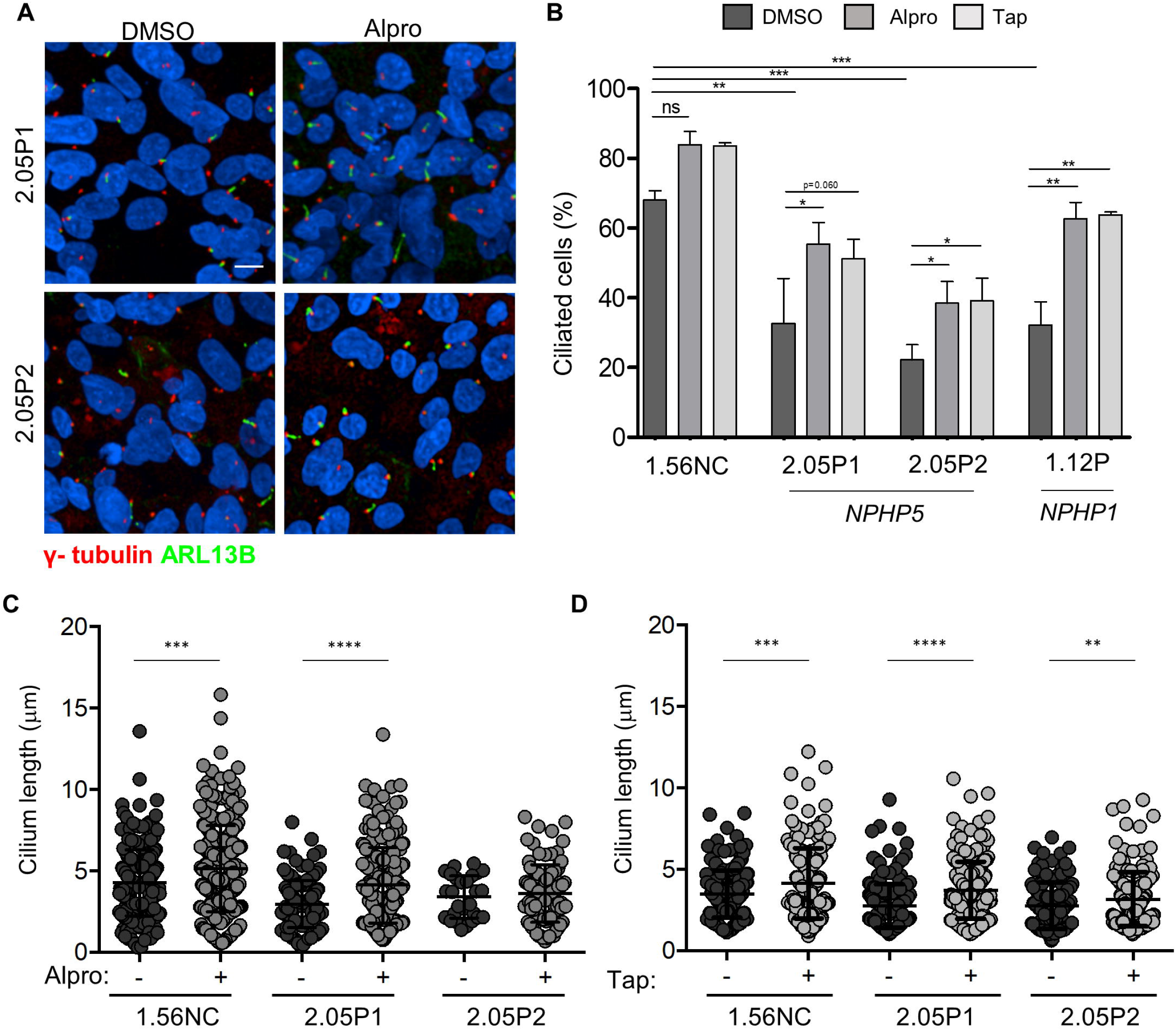
Ciliogenesis defects in *NPHP5* URECs are rescued by Prostanglandin analogues and Eupatilin treatments. (**A**) Immunofluorescence for primary cilia (ARL13B, green) and basal bodies markers (γ-tubulin, red) of *NPHP5* URECs (2.05P1, 2.05P2) treated with 0,04% DMSO or 2μM Alprostadil (Alpro) for 48h. Scale bar 10μm. (**B**) Ciliogenesis was quantified in control (1.56NC), *NPHP5* (2.05P1, 2.05P2) and *NPHP1* (1.12P) URECs treated with either DMSO (0,04%), 2μM Alprostadil (Alpro) or 1μM Tapeneprag (Tap) which were fixed and stained similarly as in **A** (n=4 experiments). Mixed linear regression with quasi-binomial penalization: \*\*\**P* < 0.001, \*\**P* < 0.01, \**P* < 0.05. (**C**) Cilium length was quantified as in Figure 1 in control (1.56NC) and *NPHP5* (2.05P1, 2.05P2) URECs treated with 0,04% DMSO (-) or either 2μM Alprostadil (Alpro, **C**) or 1μM Tapeneprag (Tap, **D**) (n=4) experiments; paired two-tailed Student’s t-test: \*\*\*\**P* < 0.0001, *** *P* < 0.001, ** *P* < 0.01.

**Figure 5:**
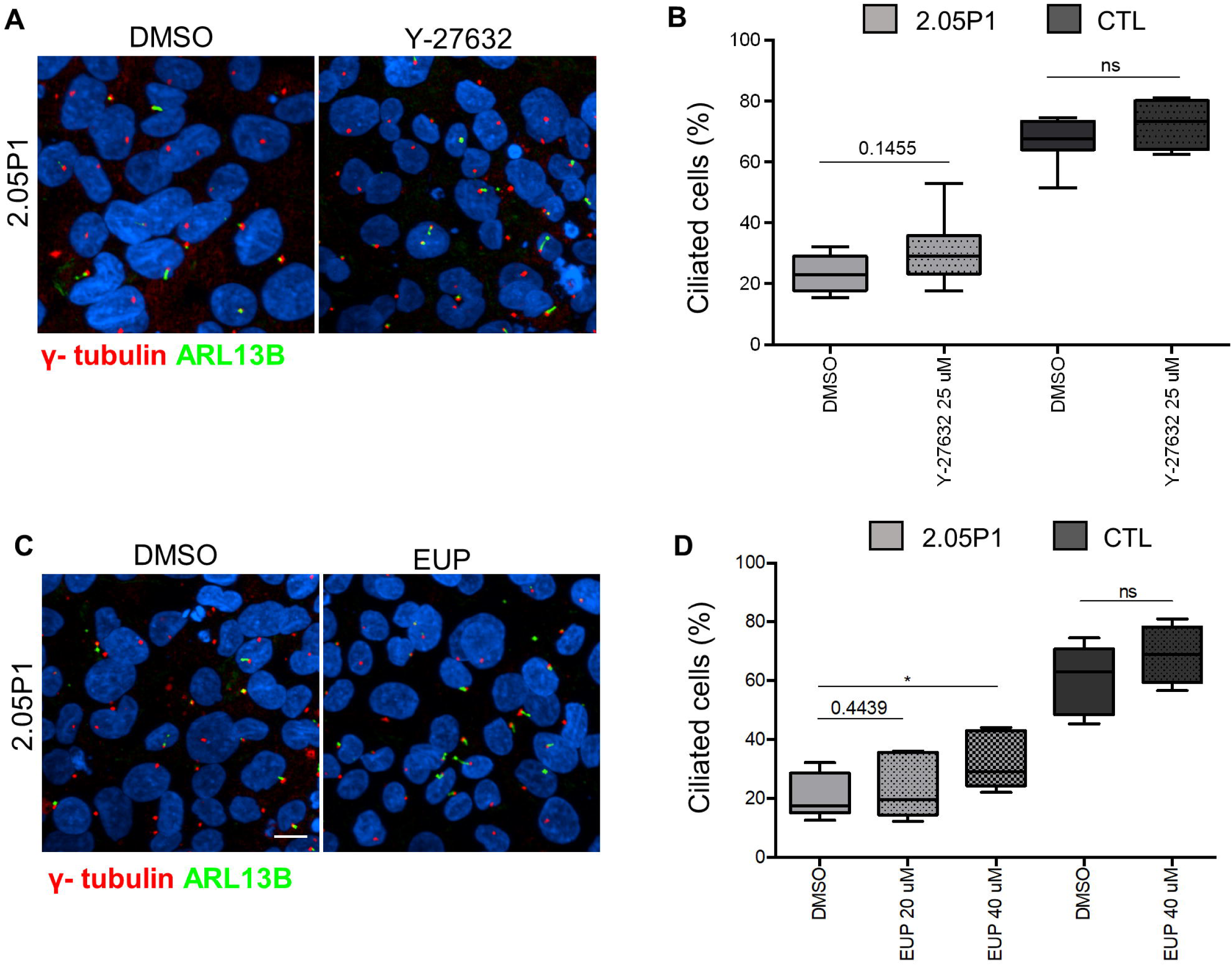
Eupatilin but not Rock inhibitor increases ciliation in *NPHP5* URECs. (**A-D**) Control (1.56NC) and *NPHP5* (2.05P1) URECs treated with DMSO (0,04%) or either ROCK inhibitor (Y-27632i, 25μM; **A** and **B**) or Eupatilin (EUP, 20μM and/or 40μM; **C** and **D**) were fixed and stained primary cilia (ARL13B, green) and basal body markers (γ-tubulin, red; **A** and **C**). Ciliogenesis was quantified similarly as in Figure 1 (EUP, **B**; Y-27632, **D**). (**B**) n=3 to 5 experiments, ** *P* < 0.05. (**D**) n=2 to 4 experiments. Statistic has been made with R script for mixed linear-regression model with quasibinomial penalization.

Altogether, these results show that prostaglandin receptor agonists and Eupatilin partially rescue ciliogenesis in the context of *NPHP5* and stress the potential wide therapeutic use of those molecules in the context of juvenile NPH.

### Prostaglandin but not Eupatilin rescues pronephric cysts in a NPH zebrafish model

To investigate the effects of Eupatilin and prostaglandin receptor agonists an *in vivo* model for NPH, we decided to use zebrafish embryos which allow rapid testing of compounds. Zebrafish was widely used to validate *NPHP* gene candidates and to investigate the effects of identified variants with characteristic ciliopathy phenotypes including body axis curvature and cysts in the proximal part of the pronephros^3^. Interestingly, a *traf3ip1* mutant line (*m649*) with a variant similar to one identified in a NPH patient - Ile17Asn vs Ile17Ser - was reported but not fully phenotypically characterized^9,33^.

To facilitate the study of its impact on kidneys, the *m649* mutation was transferred in the *Tg(wt1b:GFP)* transgenic background in which GFP is expressed in the proximal part of the pronephros^10^. Mutant embryos at 48 hours post fertilization (hpf) presented with a typical ventral curvature of the body axis with the expected proportion (∼ 25%, Figure 6A). Pronephros were analyzed by confocal microscopy in 48 hpf embryos (see Methods). As expected, pronephric cysts were only observed in curved fish (Figure 6B,C) with a drastic increase of the glomerular area (Figure 6D), correlated with a loss of PC (AcTub, Figure 6E,F). Interestingly, in contrast to what has been established in cystic mouse models^34^, pronephric cysts were not associated with increased number of mitotic cells (PH3, Figure 6E,G).

**Figure 6:**
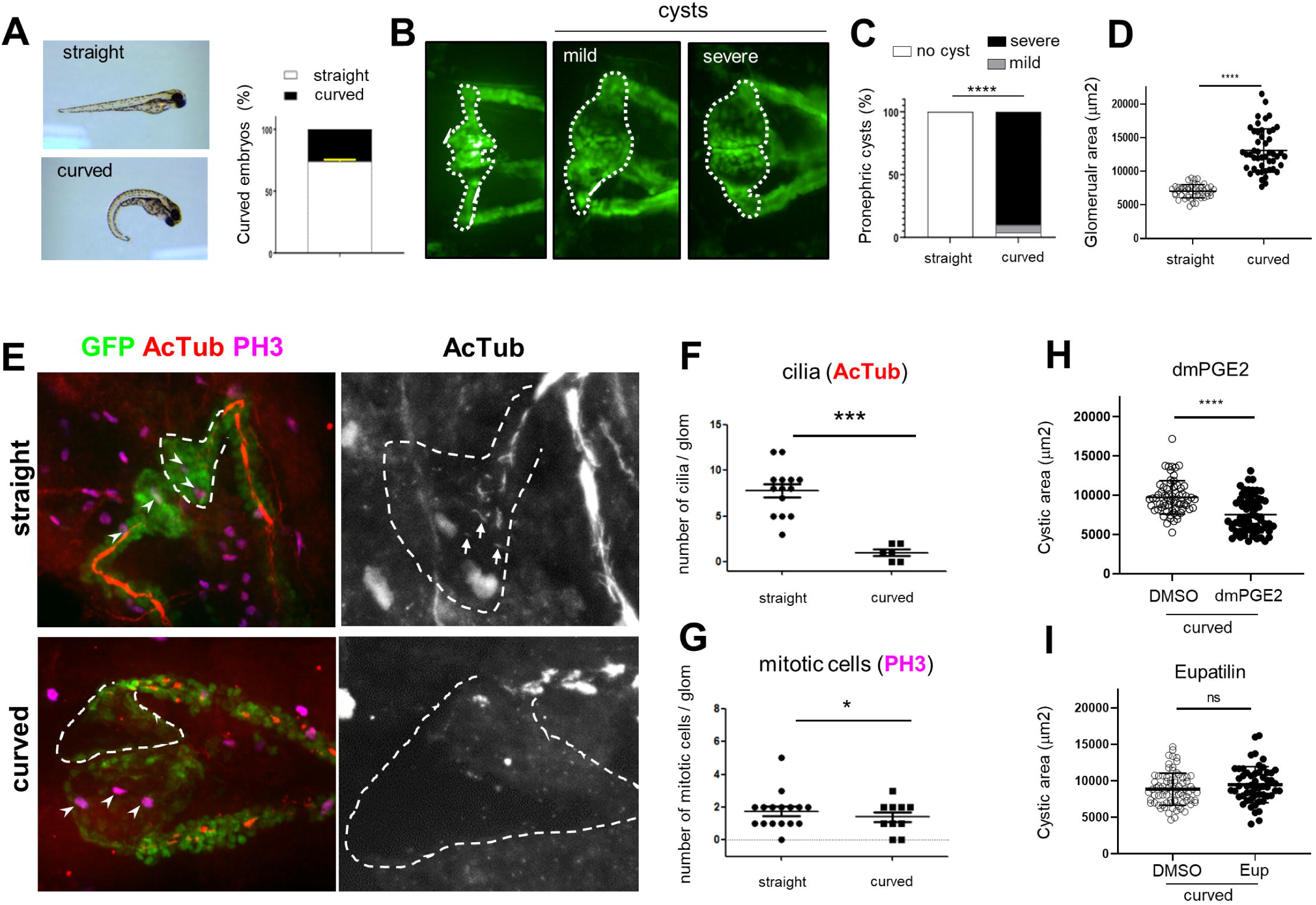
PGE2, but not Eupatilin, shows beneficial effect on cysts size in *traf3ip1* mutant fish. **(A)** Representative pictures obtained after crossing m649-GFP fish (48 hpf) with an example of straight (up) and curved (down) embryos. The curvature phenotype was quantified (n=8 experiments). **(B)** Representative pictures of the GFP-expressing pronephros with, from left to right: wild-type, mild (unilateral or small) or severe (bilateral) cystic pronephros. The region of the pronephros used to quantify cyst area is indicated by a dotted white line. **(C)** Quantification of cyst severity score in straight and curved embryos (n=4 experiments, Chi-square test: ****P<0.0001). **(D)** Quantification of cyst area measured as explained in **B** for straight and curved embryos (n=4 experiments; unpaired Student’s t test with Welch’s correction: ****P<0.0001; error bars representing mean +/- SD. **(E**) Straight (upper panels) and curved (lower panels) m649-GFP embryos were fixed and stained for cilia (acetylated-αtubulin, AcTub; red) and mitotic cells (phospho-histone-H3, PH3; purple). Representative color images are shown on the left. The areas surrounded by dotted lines represents one side of the glomerulus with the initial part of the proximal tubule based on GFP staining. On the right, zoomed (x 2.33) views of the same samples are shown for AcTub only. White arrowheads on left and arrows on right pictures indicate mitotic cells and cilia, respectively. Scale bars: 20µm. **(F)** Quantification of cilia number per glomerulus for straight and curved embryos. Unpaired Student’s t test with Welch’s correction: ****P<0.0001. **(G)** Quantification of mitotic cells number per glomerular region for straight and curved embryos. Mann-Whitney test: *P<0.05. **(H, I)** m649-GFP embryos were treated with DMSO, dimethyl-PGE2 (dmPGE2; 100 µM) or Eupatilin (2,5µM) starting at 24hpf. Quantification of the effects of treatments on cyst area (dmPGE2: Mann-Whitney test; ****P<0.0001; Eupatilin: unpaired Student’s t test with Welch’s correction: ns P>0.05).

We then tested both Eupatilin and prostaglandins in this validated NPH model. Treatment of mutant embryos with 100μM dmPGE2 for 24 hours did not rescue body axis curvature but significantly decreased cystic area in curved mutant fish (Figure 6H). As Eupatilin showed toxic effects on development when used at 40μM (most efficient concentration in URECs, see Figure 1), several concentrations were further tested starting at 1μM at 24 hpf for 24 hours. Cardiovascular defects (oedema, hemoragies) as well as pronephros migration delay were observed at, or above 5μM (supplementary Figure S8). However, when used at 2.5μM, Eupatilin did not show beneficial effects on both body axis curvature and cyst size (Figure 6I).

## Discussion

Juvenile NPH is the most common form of NPH for which, despite recent advancements, treatments remain limited to palliative care and replacement therapies. Here, we highlight the potential interest of prostaglandin analogues and Eupatilin. Our results show the broad effects of those molecules on ciliogenesis in distinct *NPHP* contexts and provide further evidence for the robust effects of PGE2 using a zebrafish NPH model. They however failed to demonstrate any positive effect of a list of potential interesting molecules from our initial screen^7^.

The first class of selected molecules was modulators of cAMP signaling, a pathway known for its positive effects on ciliogenesis^35,36^, and that we demonstrated to be critical for the PGE1-mediated effects^7^. Isoproterenol is an agonist of β2-adrenergic receptor which is implicated in cAMP production. The adrenergic system has several crucial roles in the kidney physiology but the expression of the different receptors is not homogeneous along the tubule^37^. We then hypothesized that the lack of effect of Isoproterenol in URECs could be due to the lack or poor expression of the adrenergic receptors in those cells. Theophylline, an inhibitor of phosphodiesterases^38^, was expected to increase cAMP through inhibition of its degradation. Its lack of effect is likely due to lower levels of cAMP in the absence of a specific stimulation of this pathway.

Another promising class of compounds investigated in our study were HMG-CoA reductases inhibitors, including Fluvastatin and Simvastatin, which are widely used to decrease cellular cholesterol pool. Interestingly, increased cholesterol was shown to impact cilia^39,40^. Since statins did not show positive effects on ciliogenesis under standrad UREC culture conditions with serum, a source of exogenous cholesterol, we also tested those compounds in URECs grown without serum. In such conditions, URECs showed different shape and detached from the plates which might have introduced bias into our analysis.

Finally, we tested two types of molecules which were shown to increase ciliogenesis in cancer cells in a similar phenotypic screening approach as ours^41^. The glucocorticoid dexamethasone, previously shown to modulate cilium elongation, Hh and cAMP signaling^42,43^, did not show positive effect on ciliation in URECs. This is likely due to the fact that the medium used to grow URECs contains glucocorticoids which may have masked any effect of similar molecules. In addition, among antibiotics of the cephalosporin family, neither Cefotetan nor Cefaclor showed significant effects on ciliation in URECs.

In conclusion, despite the fact that several studies indicated that the hits that we selected could be of potential interest, none of theme was able to increase ciliation in *NPHP1* URECs. It should be noticed that the screens made by other groups and ours were performed in cancer or non-human kidney cell lines grown in different media, which may then respond differently to the identified molecules compared to URECs.

RHO GTPase signaling leads to the activation of ROCK which controls actin-based cytoskeleton dynamics and contractility^44^ that negatively impacts ciliogenesis leading to decreased ciliation and/or shorter cilia^18^. Increased RHO activity has been further observed in different ciliopathy conditions where ROCK inhibitors showed positive impact on cilia *in vitro*^45–49^ as well as on kidney phenotypes in mouse models^47–49^. Interestingly, the ROCK inhibitor Y-27632 was shown to partially rescue ciliogenesis in *NPHP1* URECs, confirming the contribution of dysregulated RHOA signaling in *NPHP1*-associated phenotypes^7^. It was not efficient in *NPHP5* URECs in which RHO signaling defects remain to be investigated.

Eupatilin turned to be the most efficient among the tested molecules, showing a positive impact on ciliation in both *NPHP1* and *NPH5* URECs. This is consistent with its previously observed similar effects in the context of *NPHP6* and *NPHP8*^16,17,50^. Our transcriptomic analysis provides additional evidence for its beneficial effects on ciliogenesis. Indeed, Eupatilin mostly downregulates genes related to cell cycle progression and increases the ones required for entering into quiescence, the first step required to the cells to form PC^51^. Interestingly, Alprostadil treatment also leads to increased expression of p27^kip^^1^ a positive regulator of quiescence^7^. Surprisingly, Eupatilin was not able to modulate cyst size in *traf3ip1* mutant zebrafish embryos. This lack of effect is likely due to the fact that pronephric cysts are not associated with increased proliferation. In addition, and similarly to PGE1, RHO signaling was also downregulated upon Eupatilin treatment, which likely positively contributes to its positive effects on ciliogenesis. Altogether, our results suggest a possible convergence in the mechanism of action of these two unrelated molecules.

Finally, another aspect of our investigation focused on the study of the effects of prostaglandin-related molecules, including Alprotadil/PGE1 and Tapeneprag a specific agonist of EP2, the main PGE2 receptor expressed in URECs^7^. Both EP agonists rescued ciliogenesis in *NPHP1* URECs as well as in two *NPHP5* UREC lines.

In conclusion, similarly, as observed for Eupatilin, the present results extend our previous observations in the context of *NPHP1* and broaden the potential use of agonists of the prostaglandin receptor agonists across various NPHP contexts.

## Materials and Methods

### URECs

Patient and control individuals were recruited either at Necker or Montpellier Hospital and urine samples were collected upon written informed consent and anonymized in the frame of the NPH1 protocol approved by the French National Committee for the Protection of Persons under the ID-RCB no.2016-A00541-50. Urinary renal epithelial cells (URECs) were isolated from urine from NPH patients carrying biallelic variants either in *NPHP1* or *NPHP5* and healthy age matched donors. Control and *NPHP1* URECs were described previously^7^. All these cell lines belong to the Imagine Biocollection (declared to the French Minister of Research under the number DC-2024-6350). Briefly, *NPHP5* URECS were immortalized by retroviral transduction of thermosensitive SV40 T-antigen (LOX-CW-CRE, Addgene) at a multiplicity of infection of 5 with 8 ug/ml polybrene reagent and cells were subsequently grown in T75 flasks at the permissive temperature of 33 °C and 5% CO_2_ for maintenance, in UREC culture medium (REBM Basal Medium [CC-3139, Lonza], 1x REGM Single Quots [CC-4127, Lonza], 2% FBS certified [Invitrogen]).

### Zebrafish

The zebrafish *traf3ip1* mutant line m649^9^, a kind gift of Jarmia Malicki, was maintained at 28.5°C under standard conditions and according to European current legislation. Heterozygous m649 zebrafish were crossed with wild-type Tg(wt1b:GFP)^10^ to allow analysis of the proximal pronephros. The resulting line was called m649-GFP. For treatment with compounds, 10 hours post fertilization (hpf) embryos were transferred into 12-well plates (20 embryos/well) and incubated at 24 hpf for 24 hours with either Eupatilin or dmPGE2 diluted in 1% DMSO Egg-Water in presence of phenylthiourea (to block pigmentation). Phenotypic analysis was done on 48 hpf embryos obtained from in-cross of m649-GFP heterozygous fish.

### Compounds

The compounds selected and used in this study were: Alprostadil/PGE1 (2μM, 1620 R&D system); Taprenepag/MDT-110/CP544326 (1μM, Medetia); Y-27632-dihydrochloride (1683 Axon); Dexamethasone 21-acetate (D1881 Sigma); Isoproterenol hydrochloride (I6504 Sigma); Theophylline monohydrate (c5967-84-0 Santa Cruz); Cefotetan disodium (A5737 Sigma); Cefaclor (C6895 Sigma); Simvastatin (S6196 Sigma); Fluvastatin sodium hydrate (SML0038 Sigma); Eupatilin (SML1689 Sigma); dimethyl-PGE2 (dmPGE2; sc-201240B ChemCruz). They were dissolved either in dimethyl sulfoxyde (DMSO, Sigma) or in H_2_O, based on their solubility.

### Ciliogenesis and compound treatment

For ciliogenesis assays, URECs (10^5^ cells per well) were seeded in 96-well glass-bottomed plates (Sensoplates, Greiner) at 39°C, restrictive temperature in which cells stop to proliferate and ciliate. To test the selected compounds, cells were incubated at day 3 for 48 hours in the presence of each of them at the indicated concentration or with DMSO, and ciliogenesis was analyzed at day 5. For the other immunofluorescence experiments (ciliary composition), URECs (4 × 10^5^ cells per well) were seeded on glass-bottomed coverslips in 24-well plates at the same temperature as for ciliogenesis.

### Immunofluorescence

Cells were fixed in either cold methanol for 5 min or with 4% PFA for 20 min, quenched in 50 mM NH_4_Cl, and permeabilized with 0.1% Triton 15 min when fixed with 4% PFA. Incubation of primary and secondary antibodies was performed in PBS, 0.1% Tween-20, 3% BSA (Sigma-Aldrich), both 1h at room temperature. Nuclear staining was performed using 4′,6-diamidino-2-phenylindole (DAPI 1/2000e, Thermo Fisher Scientific, 62247). For ciliogenesis experiments, cells were left in PBS (96 well plates); for the other immunofluorescence, coverslips were mounted using Mowiol mounting medium on Superfrost slides (Thermo Fisher). Zebrafish embryos (48 to 50 hpf) were fixed in 4% PFA overnight at 4°C and placed in methanol 100% gradually. The embryos were removed from methanol then washed in PBS/triton 0.5%. Post-fixation was done in PFA 4% for 20 minutes. Fixed embryos were then incubated in blocking buffer (PBS, triton 0,5%, 10% FBS, 2% BSA) for at least 1 hour at room temperature. Incubation with primary and secondary antibodies were performed in blocking buffer at 4°C overnight and at 37° for 2 hours, respectively. The immunostained embryos were washed during one day in PBS on a rocker at 4°C. Fixed and stained embryos were mounted in 1% low gelling temperature agarose (Sigma-Aldrich, A9414) on µ-Dish 35mm, high Glass Bottom Ibidi (81158).

### Antibodies

The following antibodies were used: acetyl-α tubulin mouse monoclonal (1/300; Sigma, T6793), ADCY3 rabbit polyclonal (1/100e; Invitrogen, PA5-35382), ARL13B rabbit polyclonal (1/800e; ProteinTech, 17711-1-AP), ARL13B mouse monoclonal (1/100e; ABCAM, ab136648), INPP5E rabbit polyclonal (1/200e; Proteintech, 17797-1-AP), γ-Tubulin mouse monoclonal (1/5000e; Sigma, T6557), GT335 mouse monoclonal (1/5000e; Adipogen, AG-20B-0020-C100), PH3 rabbit polycolonal (1/200; Cell Signaling Technology, 3377S), NPHP4 rabbit polyclonal (1/100e; BiCell scientific, 90004), NPHP11 rabbit polyclonal (1/100e; BiCell, 90103). Secondary antibodies (donkey) conjugated to Alexa Fluor® 488, 555 or 647 were used (1/1000e; Molecular Probes, Thermo Fisher Scientific).

### Image acquisition, quantification, and analysis

For ciliogenesis assays, images were acquired using the Opera Phenix microscope (40X water, Perkin-Elmer) and automated acquisition of 41 z-stack per well was performed. The percentage of ciliated cells was measured using a semi-automated workflow with Harmony software (Perkin-Elmer). Images were analyzed as previously^7^ using the building blocks approach to detect in sequence: nuclei (DAPI staining) and, with a 20-pixel enlarged region, the basal body (γ-Tubulin staining), and then the primary cilia (ARL13B staining). With filters (intensity, size, distance to the putative basal body, signal-to-noise ratio), the software segmented candidate primary cilia. Then, multiple phenotypic parameters were calculated for every candidate cilium using signal enhancement ratio texture (intensity patterns) and advanced STAR morphology parameters. Using the PhenoLOGIC machine-learning option of Harmony, the parameters best suited to discriminate cilia were defined and used to obtain final detection and counting. For other assays with cell mounted on microscope slides, images were acquired using either a Spinning Disk microscope (40X or 63X, Zeiss) or an epi-illumination microscope (Leica, DMR) with a cooled charge-coupled device camera (DFC3000G, Leica). All quantifications were performed using Fiji (v2.14.0) opensource software^11^.

For ciliary composition analyses (ADCY3, INPP5E) it was used a semi-automated approach using Fiji coupled with Ilastik (v1.3.3post3) opensource softwares. Briefly: a first macro in Fiji splits and generates Max intensity projections files for the cilium channel (ARL13B) and for the protein of which is required to calculate the abundancy along the cilium (ADCY3, INPP5E). Images of cilium channel are then charged on Ilastik a program that uses machine-learning pixel classification to recognize true vs false signal of interest generating a simple segmentation file. A second macro uses the ilastik simple segmentation mask to generate cilium region of interest (ROI) with the possibility to check and correct it manually. The last macro in Fiji then calculates for each cilium the area and the length, plus the mean gray value and integrated density of protein of interest channel already filtered from the noise signal. Quantification of the intensity of transition zone proteins were performed using Fiji.

Cysts in the proximal pronephros were analyzed on live anesthetized 48 hpf embryos using the Opera Phenix (PerkinElmer) microscope using a 10x dry objective. GFP-positive embryos were individually positioned on the dorsal side in 96-well plates using a previously published method^12^ to perform automated imaging of the proximal pronephros with Opera Phenix HSC system (Perkin Elmer). Percentage of embryos with pronephric cysts was quantified manually (severe: bilateral big cysts; mild: unilateral or small bilateral cysts) or using Fiji to measure area of cysts with manual selection of the region corresponding to the glomerulus and neck regions of the pronephros. Fixed and immunostained embryos were imaged using a Zeiss Axio Observer Z1 inverted microscope equipped with a Yokogawa CSU-X1 spinning disk. Images were acquired with a 20× dry objective through a Hamamatsu Orca Flash 4.0 sCMOS camera.

### RNA extraction and RNA-seq analysis

URECs were seeded in 12-well plates at 39°C grown for 3 days and then incubated for 24 hours with Eupatilin (20μM) or DMS0 (0,20%). Cells were lysed in RLT buffer and mRNA was isolated using an Extraction Mini Kit (Qiagen) following the recommendations of the manufacturer. cDNA library was prepared using the Universal Plus mRNA stranded kit (Tecan-Nugen). Then, for each sample, 50-million passing filter reads/clusters (paired-end 100+100 bases) was produced on NovaSeq6000 Illumina. FASTQ files were mapped to the ENSEMBL [Human(GRCh38/hg38)] reference using Hisat2 and counted by featureCounts from the Subread R package (http://www.r-project.org/). Read-count normalizations and group comparisons were performed by the Deseq2 statistical method^13^. The results were filtered at P ≤0.05 and fold-change 1.2. Heatmaps were made with the R package ctc: Cluster and TreeConversion, and imaged by Java Treeview software (Java Treeview). Functional analyses were carried out using Metascape (http://metascape.org/).

### qRT-PCR Analysis

Total RNA was reverse-transcribed using SuperScript II Reverse Transcriptase (LifeTechnologies) according to the manufacturer’s protocol. Quantitative real-time PCR (qRT-PCR) was performed with iTaq Universal SYBR Green Supermix (Bio-Rad) on the CFX-384 Real-Time PCR System (Biorad). Each biological replicate was measured in technical duplicates. Primers used for qRT-PCR are listed in Supplementary Table 2. For quatification and normalization it was used the method of multiple control genes, previously decribed^14,15^, step 7.

### Sanger Sequencing

Total DNA was isolated from *NPHP5* URECs using the NucleoSpin Mini Kit (Machery-Nalgene) following the recommendations of the manufacturer. PCR was performed with Mytaq (Neb/Biolabs). For sequencing reactions were used: Exosap (Illustra), Big-dye (Thermo-Fisher) according to the manufacturer’s protocol. Primers: Fw-exon6-CTGTATCCTCTAACTCCCCAGA; Rev-exon6-TGTGGTTATCTGCACAGTTCA; Fw-exon7-CAGAAAGTTTAGCAGAGATGGTC; Rev-exon7-TGAAGTTCCATCACCATTGT. Genotyping of zebrafish embryos was performed by placing embryos in 10 mm Tris pH 8.0, 1 mm EDTA and 1.2 mg/ml at 55°C overnight, and the reaction was stopped by incubating at 95°C for 5 min. DNA from exon 1 was Sanger sequenced using the following primers: forward 5′GAA-CGA-ATC-GGT-AGC-CAA-AA3′ and reverse 5′AGA-CTA-CGC-GCA-GGC-TAC-AT3′. Sequencing was performed on 3500xL Genetic Analyzer (Thermo Fisher) and analyzed using Sequencher software (Gene Codes).

### Statistics

Statistical calculations were performed with R or GraphPad Prism v6. Figure representation was performed using GraphPad Prism v6 and is presented as dot plots, box-and-whisker plots or bar plots. Differences between the same group meeting the normal distribution were analyzed using paired two-tailed Student’s t-test; if not possible or if data do not meet the normal distribution data were compared using the Wilcoxon test. Differences between two groups meeting the normal distribution were compared using unpaired two-tailed Student’s t test. To analyze RT-qPCR data after Eupatilin treatment in cells coming from the same individuals, we used a paired two-tailed Student’s t test. To analyze the effects of different compound treatments on ciliogenesis in the same cell line, a mixed linear-regression model with quasi-binomial penalization (R software: qlmmPQL function of MASS package) was used, taking into account the correlation of observations coming from the same individuals using a random effect on the cell line. Statistical tests and exact sample sizes used to calculate statistical significance are stated in the figure legends.

## Acknowledgment

AB and SS were supported by grants from the Agence Nationale de la Recherche (ANR), including the “Investissements d’Avenir” program (ANR-10-IAHU-01) and the “RHU-C’IL-LICO” as part of the second “Investissements d’Avenir” program (reference: ANR-17-RHUS-0002). AT and GR were supported by a fellowship from European Commission Horizon 2020 research and innovation programme Marie Sklodowska-Curie Innovative Training Networks (grant number: 861329) and by the TheRaCil consortium (Horizon-health-2022-disease-06-two stage, grant 101080717), respectively. MH was supported by ORKID. We thank the patients and their families for their participation, the LEAT and cell imaging facilities of the SFR Necker (Structure Fédérative de Recherche; Inserm US24, CNRS UAR 3633), as well as Amandine Viau for her help on RNAseq and qPCR analyses.

## Authors’ roles

AB designed and supervised the study. MF, EP and SS collected the family samples and arranged clinical testing. AT, EP, AS, SS, and AB designed cell biology in vitro experiments which were performed and analyzed by AT. GR, AT, MH and AB designed cell biology zebrafish experiments which were done and analyzed by MH and GR. NG and AT settled up the macro for cilia analysis by immunofluorescence and AT analyzed the data. CB and NC generated and analyzed RNAseq data with AT. JPA and LBR provided prostaglandin-related molecules and helped with image analyses. AT and AB wrote the manuscript.

## Disclaimers

JPA and LBR are shareholders at Medetia Pharmaceuticals; SS, LBR, JPA are authors in the patent application WO2109/075369A1, currently on PCT National examination phase. There are no other conflicts of interest.

