## supplementary data for "Evaluation of Prostaglandin Receptor Agonists and Eupatilin in the Context of Nephronophthisis"

**Supplementary Figure legends**

**Figure S1: Hits from the Prestwick library screening in *NPHP1* URECs.**
**Figure S2: Cell cycle genes modulated upon EUP treatment.**
**Figure S3: RHO and Autophagy gene sets modulated upon EUP treatment.**
**Figure S4: Ciliary gene set modulated upon EUP treatment.**
**Figure S5: Genes commonly modulated by EUP and ALP treatments.**
**Figure S6: Characterization of URECs from two individuals with heterozygous compound variants in *NPHP5***
**Figure S7: *NPHP5* URECs show ciliogenesis and ciliary composition defects.**
**Figure S8: Determination of nontoxic dose of Eupatilin for treatment of zebrafish embryos**

**Supplementary Table 1**

**Supplementary Figure legends**

**Figure S1: Hits from the Prestwick library screening in *NPHP1* URECs.**

**(A)** Control (1-56NC) and *NPHP1* (1-12P, 1.03P) URECs grown in ciliogenesis conditions for 5 days at non permissive temperature (39°C) were fixed and stained for primary cilia (ARL13B, green) and basal bodies (γ- tubulin, red) to score for ciliogenesis after compound treatment at day 3 for 48 hours in complete medium except for Statins. **(A')** Same dataset as in **A**, showing ciliated cells as ratio over DMSO condition after compound treatment. All experiments are n=2. Statistic test explained in Methods.

**Figure S2: Cell cycle genes modulated upon EUP treatment.**

**(A)** Heatmap showing differential expression of cell-cycle G1/S specific genes from RNA-seq dataset of *NPHP1* URECs DMSO versus EUP. **(B)** Heatmap showing differential expression of cell-cycle G2/M specific genes from RNA-seq dataset of *NPHP1* URECs DMSO versus EUP. **(C)** Heatmap showing differential expression of cell-cycle G0/G1 specific genes from RNA-seq dataset of *NPHP1* URECs DMSO versus EUP

**Figure S3: RHO and Autophagy gene sets modulated upon EUP treatment.**

**(A)** Heatmap showing differential expression of Rho/actin specific genes from RNA-seq dataset of *NPHP1* URECs DMSO versus EUP. **(B)** Heatmap showing differential expression of autophagy specific genes from RNA-seq dataset of *NPHP1* URECs DMSO versus EUP.

**Figure S4: Ciliary gene set modulated upon EUP treatment.**

Heatmap showing differential expression of Ciliary specific genes from RNA-seq dataset of *NPHP1* URECs DMSO versus EUP (An exhaustive list is on File S4).

**Figure S5: Genes commonly modulated by EUP and ALP treatments.**

**(A)** Venn diagram showing genes modulated by EUP and ALP treatments. **(B)** Heatmap showing differential expression of genes in common with ALP treatment, from RNA-seq dataset of *NPHP1* URECs DMSO versus EUP (An exhaustive list is on File S4).

**Figure S6: Characterization of URECs from two individuals with heterozygous compound variants in *NPHP5***

**(A)** Schematic representation of IQCB1/NPHP5 protein domains and exons with the position of the variants identified in 2.05P1 and 2.05P2 affected individuals. **(B)** *NPHP5* variants identified in in 2.05P1 and 2.05P2 patients was validated in URECs by Sanger sequencing (boxed in red). **(C)** Relative expression of *NPHP5* mRNA was quantified by qRT-PCR using primers specific for two different exons 4-5 and exons 13-14 junction regions. Unpaired t-test $^{**}P < 0.01$, $^{***}P < 0.001$.

**Figure S7: *NPHP5* URECs show ciliogenesis and ciliary composition defects.**

**(A)** Ciliated control (1.56.NC) and *NPHP5* (2.05P1) were fixed and stained for basal body and axoneme (GT335, red) and for ADCY3 (green). **(B)** Intensity of ADCY3 staining in cilia was quantified as explained in Methods. n=3; unpaired t test, $^{**}P < 0.01$, Scale bars 10μm. **(C)** Positive cells for ADCY3 staining in cilia was quantified as explained in Methods. n=3; unpaired Student's t test, $^{**}P < 0.01$, $^{***}P < 0.001$. **(D)** Ciliated control (1.56.NC) and *NPHP5* (2.05P1) fixed and stained for basal body and axome (GT335, red) and for NPHP4 (green) which is a marker of the transition zone (arrows). **(E)** NPHP4 staining intensity at TZ was quantified as detailed in the Methods, n=2.

**Figure S8: Determination of nontoxic dose of Eupatilin for treatment of zebrafish embryos**

(**A-C**) Wild type *Tg(wt1b:GFP)* zebrafish embryos were treated with Eupatilin (Eup) at the indicated concentrations for 24 hrs starting at 24 hpf. (**A**) The impact of Eupatilin on body axis morphology was quantified as percentage of embryos presenting with dorsal, ventral or lateral curvature. (**B**) Proximal pronephros was imaged thanks to GFP expression and the effect of Eupatilin treatment on morphogenesis was quantified and expressed as percentage of abnormal pronephros. (**C**) The presence of edema and/or yolk misorganization (arrows) was quantified and expressed as the percentage of abnormal embryos.

**Figure S1:**

A

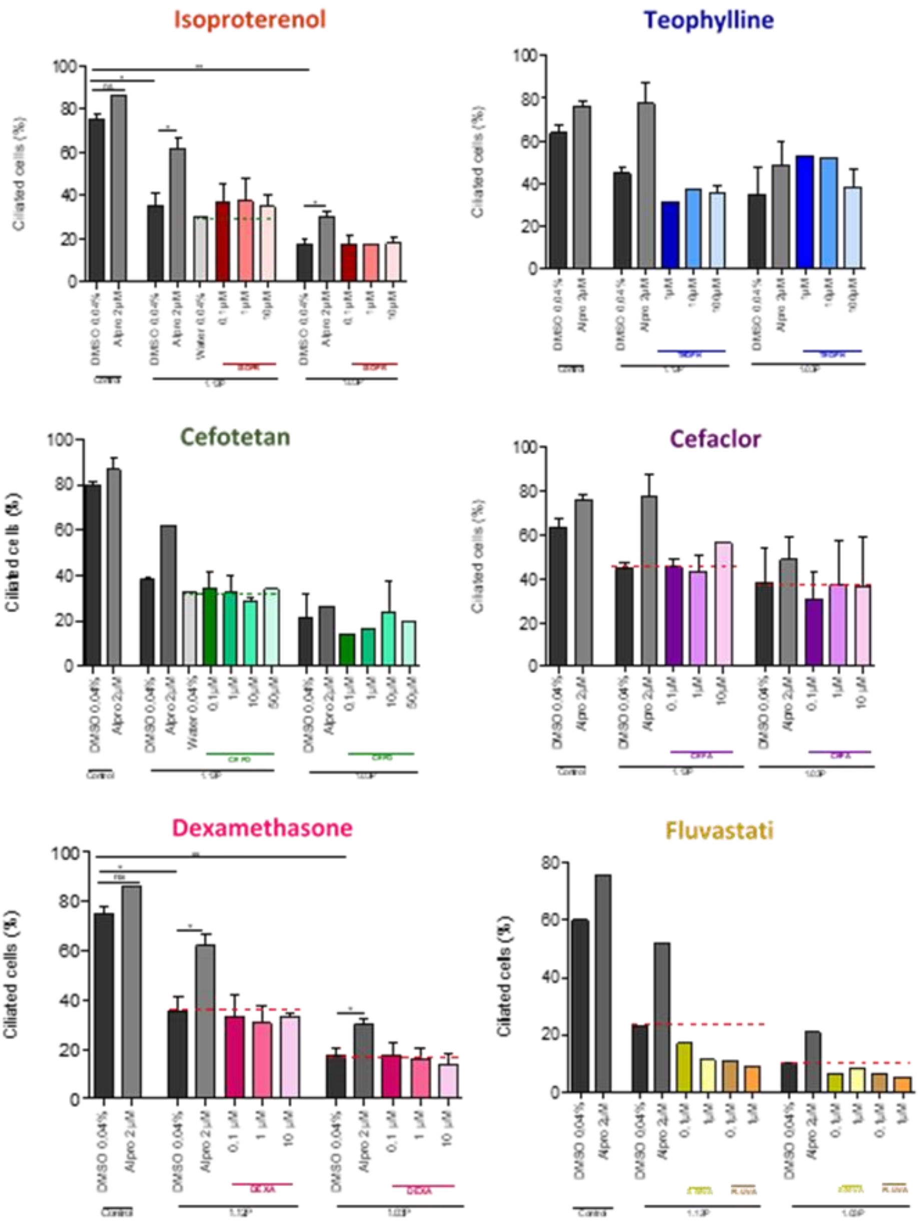
Isoproterenol
Teophylline
Cefotetan
Cefaclor
Dexamethasone
Fluvastati
Ciliated cells (%)
DMSO 0.04%
Alpro 2µM
Water 0.04%
0.1µM
1µM
10µM
50µM

A

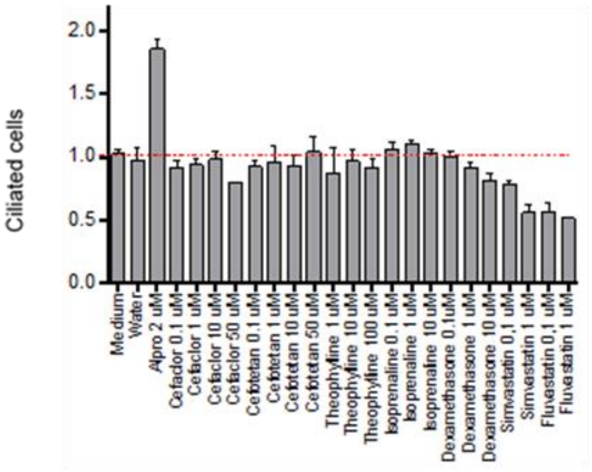
Ciliated cells
2.0
1.5
1.0
0.5
0.0
Medium
Water
Alpro 2 uM
Cefador 0.1 uM
Cefaclor 1 uM
Cefaclor 10 uM
Cefaclor 50 uM
Cefotetan 0.1 uM
Cefotetan 1 uM
Cefotetan 10 uM
Cefotetan 50 uM
Theophylline 1 uM
Theophylline 10 uM
Theophylline 100 uM
Isoprenaline 0.1 uM
Isoprenaline 1 uM
Isoprenaline 10 uM
Dexamethasone 0.1uM
Dexamethasone 1 uM
Dexamethasone 10 uM
Simvastatin 0,1 uM
Simvastatin 1 uM
Fluvastatin 0,1 uM
Fluvastatin 1 uM

**Figure S2:**

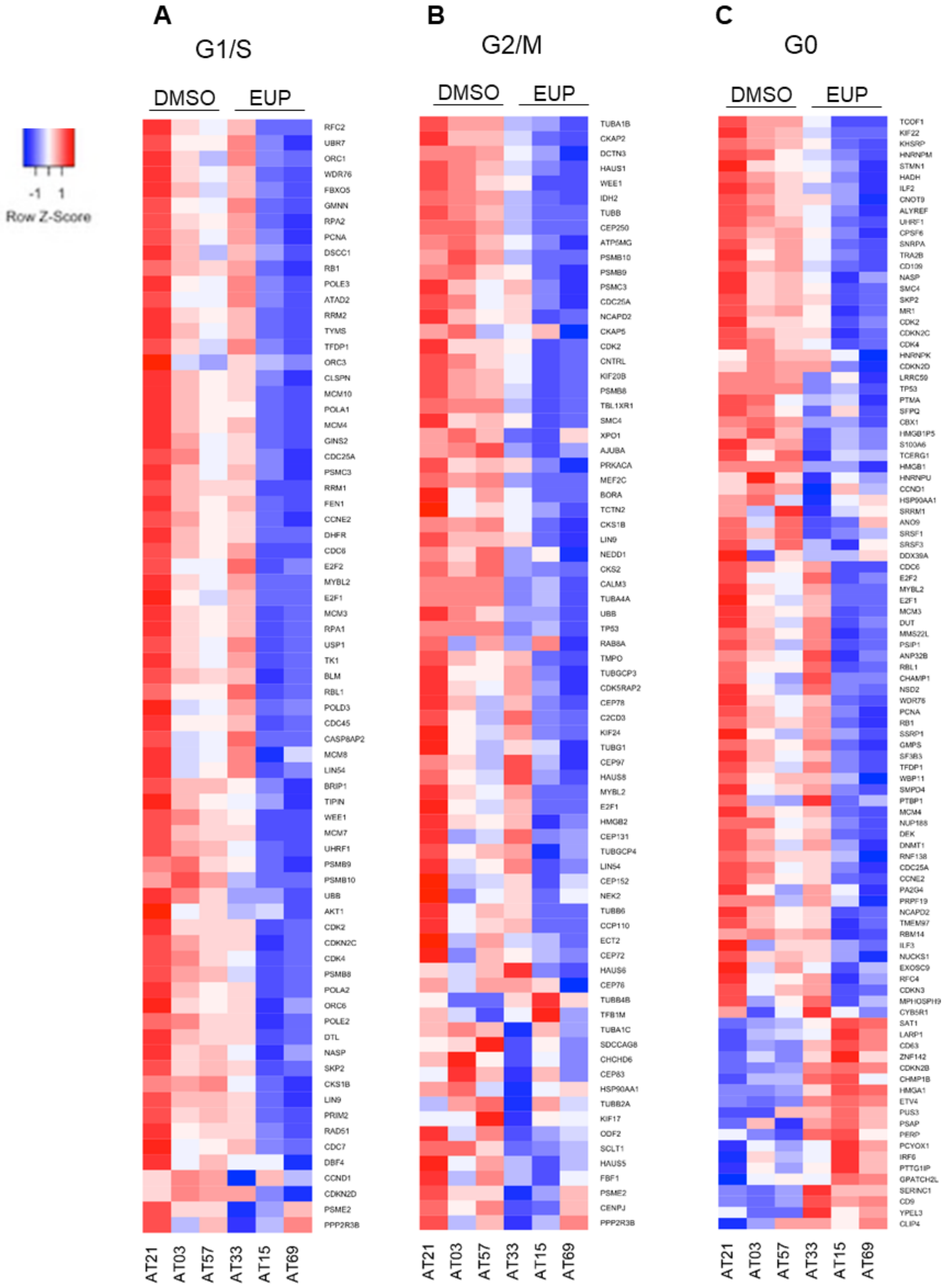
-1 1
Row Z-Score
A
G1/S
DMSO
EUP
RFC2
UBR7
ORC1
WDR76
FBXO5
GMNN
RPA2
PCNA
DSCC1
RB1
POLE3
ATAD2
RRM2
TYMS
TFDP1
ORC3
CLSPN
MCM10
POLA1
MCM4
GINS2
CDC25A
PSMC3
RRM1
FEN1
CCNE2
DHFR
CDC6
E2F2
MYBL2
E2F1
MCM3
RPA1
USP1
TK1
BLM
RBL1
POLD3
CDC45
CASP8AP2
MCM8
LIN54
BRIP1
TIPIN
WEE1
MCM7
UHRF1
PSMB9
PSMB10
UBB
AKT1
CDK2
CDKN2C
CDK4
PSMB8
POLA2
ORC6
POLE2
DTL
NASP
SKP2
CKS1B
LIN9
PRIM2
RAD51
CDC7
DBF4
CCND1
CDKN2D
PSME2
PPP2R3B
AT21
AT03
AT57
AT33
AT15
AT69
B
G2/M
DMSO
EUP
TUBA1B
CKAP2
DCTN3
HAUS1
WEE1
IDH2
TUBB
CEP250
ATP5MG
PSMB10
PSMB9
PSMC3
CDC25A
NCAPD2
CKAP5
CDK2
CNTRL
KIF20B
PSMB8
TBL1XR1
SMC4
XPO1
AJUBA
PRKACA
MEF2C
BORA
TCTN2
CKS1B
LIN9
NEDD1
CKS2
CALM3
TUBA4A
UBB
TP53
RAB8A
TMPO
TUBGCP3
CDK5RAP2
CEP78
C2CD3
KIF24
TUBG1
CEP97
HAUS8
MYBL2
E2F1
HMGB2
CEP131
TUBGCP4
LIN54
CEP152
NEK2
TUBB6
CCP110
ECT2
CEP72
HAUS6
CEP76
TUBB4B
TFB1M
TUBA1C
SDCCAG8
CHCHD6
CEP63
HSP90AA1
TUBB2A
KIF17
ODF2
SCLT1
HAUS5
FBF1
PSME2
CENPJ
PPP2R3B
AT21
AT03
AT57
AT33
AT15
AT69
C
G0
DMSO
EUP
TCOF1
KIF22
KHSRP
HNRNPM
STMN1
HADH
ILF2
CNOT9
ALYREF
UHRF1
CPSF6
SNRPA
TRA2B
CD109
NASP
SMC4
SKP2
MR1
CDK2
CDKN2C
CDK4
HNRNPK
CDKN2D
LRRC59
TP53
PTMA
SFPQ
CBX1
HMGB1P5
S100A6
TCERG1
HMGB1
HNRNPU
CCND1
HSP90AA1
SRRM1
ANO9
SRSF1
SRSF3
DDX39A
CDC6
E2F2
MYBL2
E2F1
MCM3
DUT
MMS22L
PSIP1
ANP32B
RBL1
CHAMP1
NSD2
WDR76
PCNA
RB1
SSRP1
GMPS
SF3B3
TFDP1
WBP11
SMPD4
PTBP1
MCM4
NUP188
DEK
DNMT1
RNF138
CDC25A
CCNE2
PA2G4
PRPF19
NCAPD2
TMEM97
RBM14
ILF3
NUCKS1
EXOSC9
RFC4
CDKN3
MPHOSPH9
CYB5R1
SAT1
LARP1
CD63
ZNF142
CDKN2B
CHMP1B
HMGA1
ETV4
PUS3
PSAP
PERP
PCYOX1
IRF6
PTTG1IP
GPATCH2L
SERINC1
CD9
YPEL3
CLIP4
AT21
AT03
AT57
AT33
AT15
AT69

**Figure S3:**

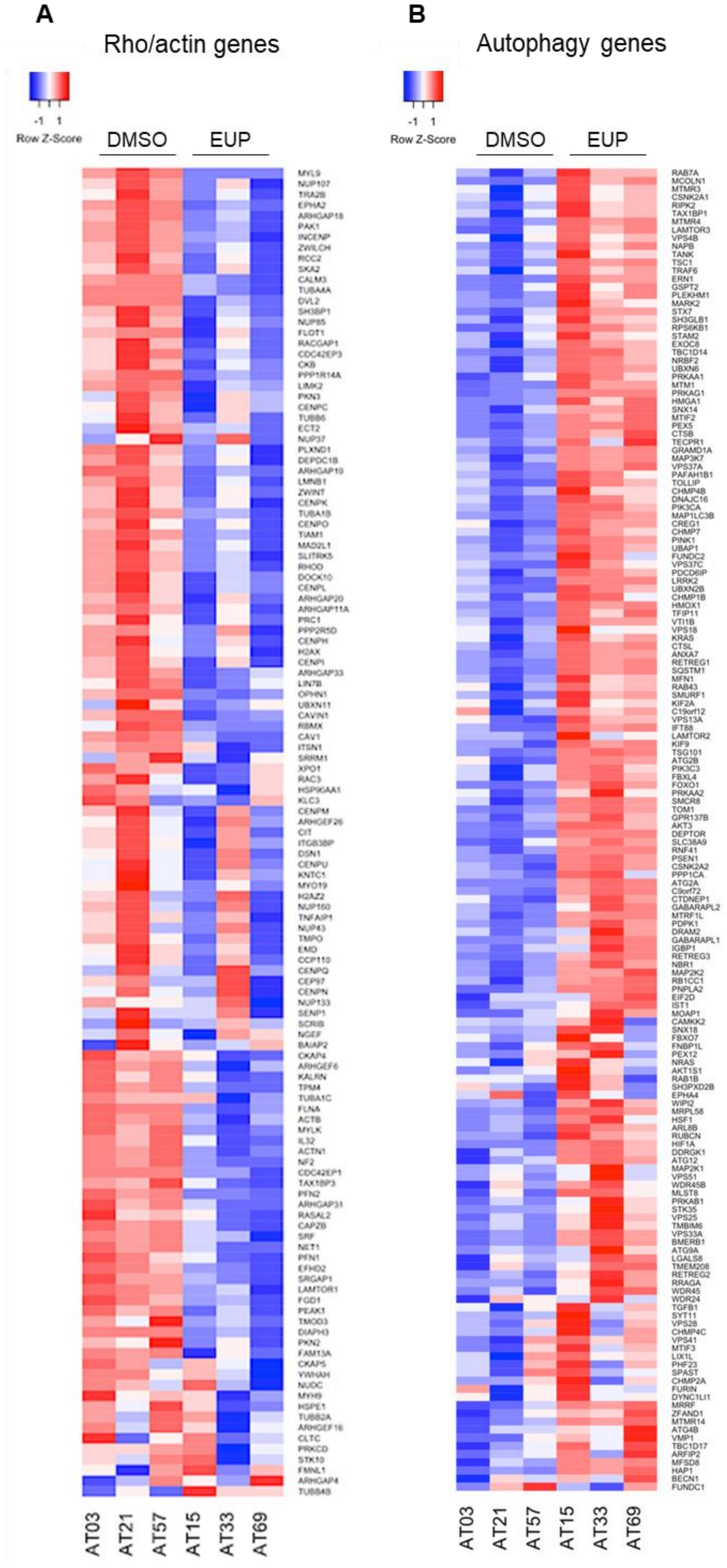
A
Rho/actin genes
-1 1
Row Z-Score
DMSO
EUP
AT03
AT21
AT57
AT15
AT33
AT69
B
Autophagy genes
-1 1
Row Z-Score
DMSO
EUP
AT03
AT21
AT57
AT15
AT33
AT69

**Figure S4:**

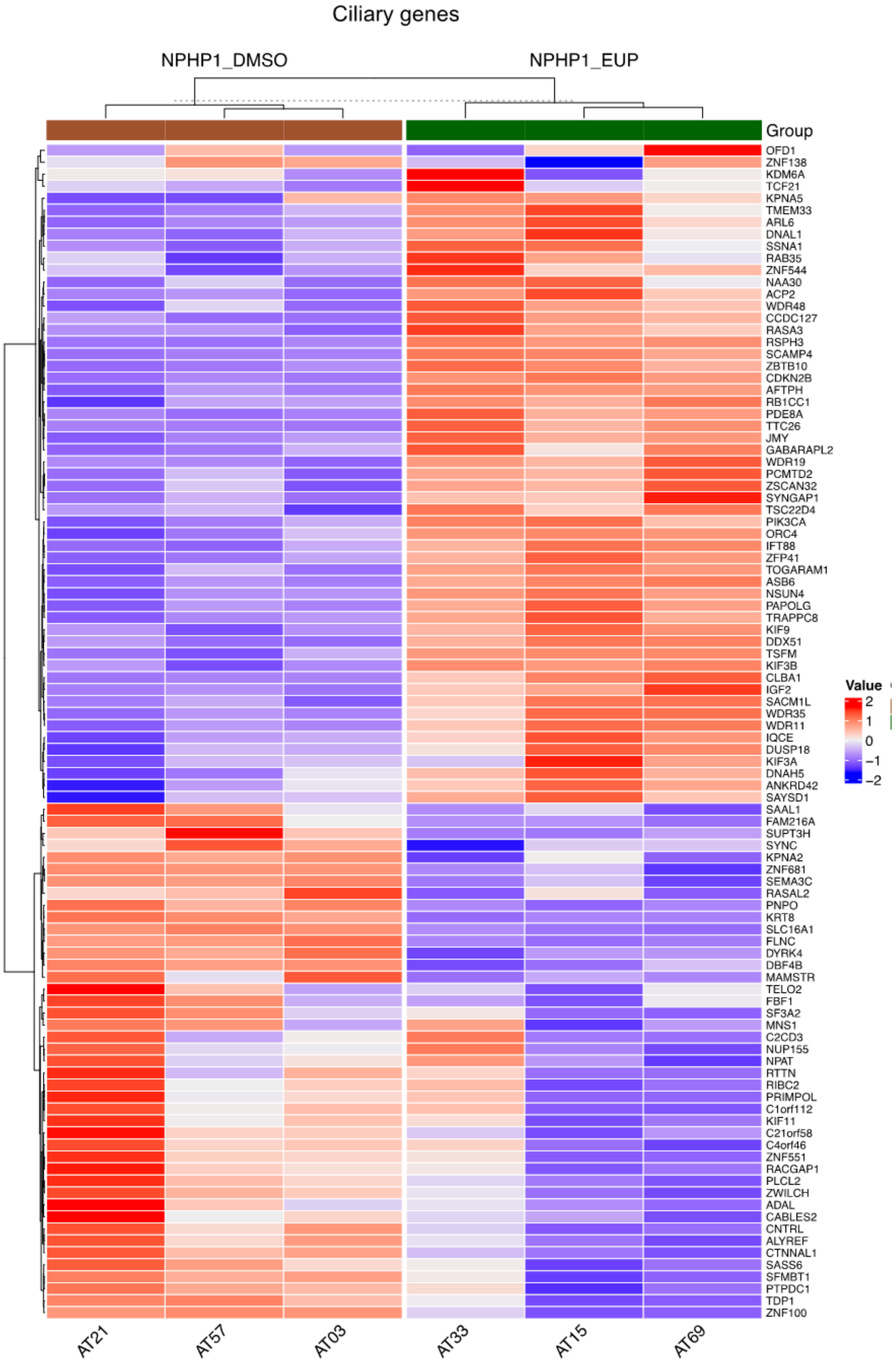
Ciliary genes
NPHP1_DMSO
NPHP1_EUP
Group
OFD1
ZNF138
KDM6A
TCF21
KPNA5
TMEM33
ARL6
DNAL1
SSNA1
RAB35
ZNF544
NAA30
ACP2
WDR48
CCDC127
RASA3
RSPH3
SCAMP4
ZBTB10
CDKN2B
AFTPH
RB1CC1
PDE8A
TTC26
JMY
GABARAPL2
WDR19
PCMTD2
ZSCAN32
SYNGAP1
TSC22D4
PIK3CA
ORC4
IFT88
ZFP41
TOGARAM1
ASB6
NSUN4
PAPOLG
TRAPPC8
KIF9
DDX51
TSFM
KIF3B
CLBA1
IGF2
SACM1L
WDR35
WDR11
IQCE
DUSP18
KIF3A
DNAH5
ANKRD42
SAYSD1
SAAL1
FAM216A
SUPT3H
SYNC
KPNA2
ZNF681
SEMA3C
RASAL2
PNPO
KRT8
SLC16A1
FLNC
DYRK4
DBF4B
MAMSTR
TELO2
FBF1
SF3A2
MNS1
C2CD3
NUP155
NPAT
RTTN
RIBC2
PRIMPOL
C1orf112
KIF11
C21orf58
C4orf46
ZNF551
RACGAP1
PLCL2
ZWILCH
ADAL
CABLES2
CNTRL
ALYREF
CTNNAL1
SASS6
SFMBT1
PTPDC1
TDP1
ZNF100
Value
2
1
0
−1
−2
AT21
AT57
AT03
AT33
AT15
AT69

**Figure S5:**

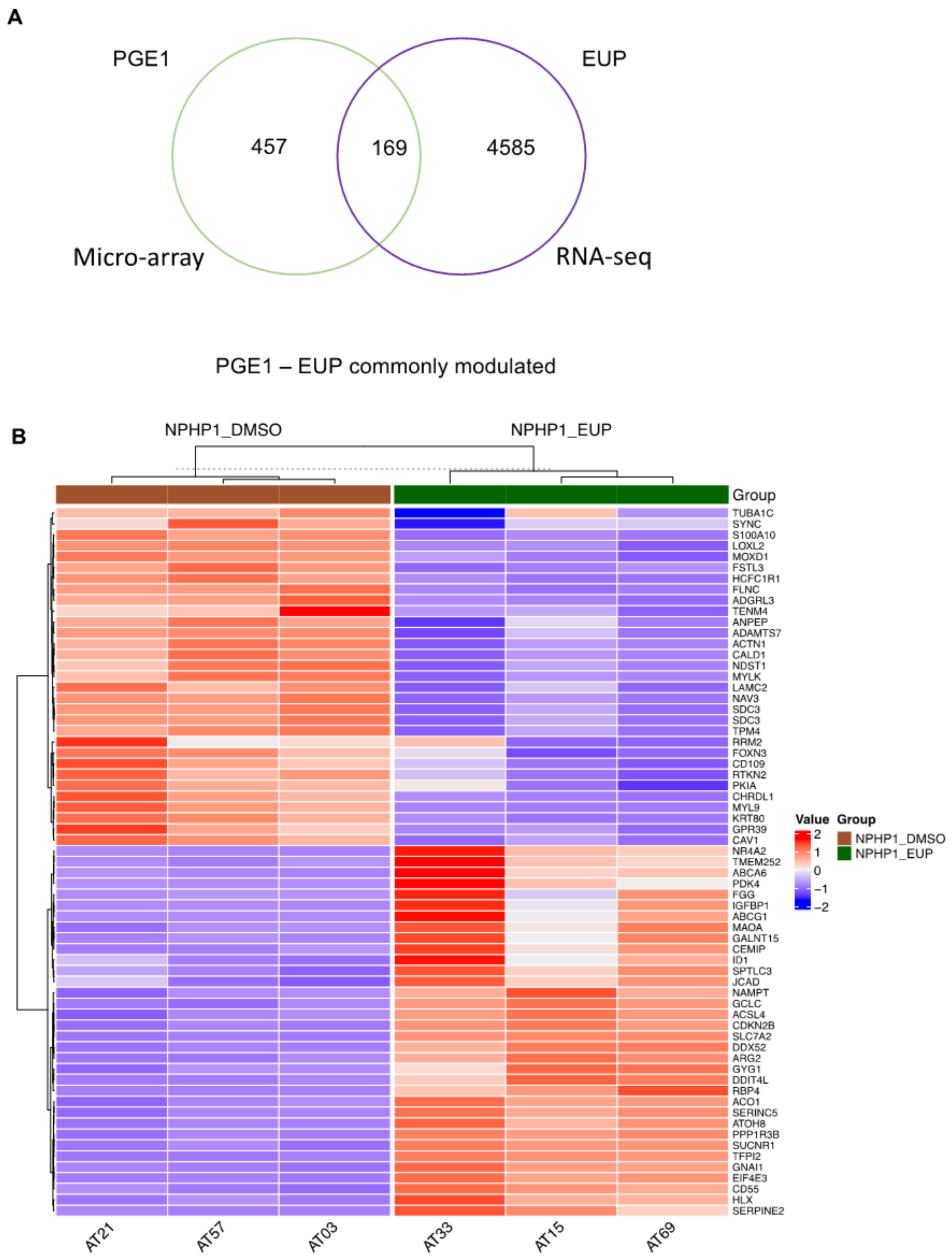
A
PGE1
EUP
457
169
4585
Micro-array
RNA-seq
PGE1 – EUP commonly modulated
B
NPHP1_DMSO
NPHP1_EUP
Group
TUBA1C
SYNC
S100A10
LOXL2
MOXD1
FSTL3
HCFC1R1
FLNC
ADGRL3
TENM4
ANPEP
ADAMTS7
ACTN1
CALD1
NDST1
MYLK
LAMC2
NAV3
SDC3
SDC3
TPM4
RRM2
FOXN3
CD109
RTKN2
PKIA
CHRDL1
MYL9
KRT80
GPR39
CAV1
NR4A2
TMEM252
ABCA6
PDK4
FGG
IGFBP1
ABCG1
MAOA
GALNT15
CEMIP
ID1
SPTLC3
JCAD
NAMPT
GCLC
ACSL4
CDKN2B
SLC7A2
DDX52
ARG2
GYG1
DDIT4L
RBP4
ACO1
SERINC5
ATOH8
PPP1R3B
SUCNR1
TFPI2
GNAI1
EIF4E3
CD55
HLX
SERPINE2
AT21
AT57
AT03
AT33
AT15
AT69
Value
2
1
0
−1
−2
Group
NPHP1_DMSO
NPHP1_EUP

**Figure S6:**

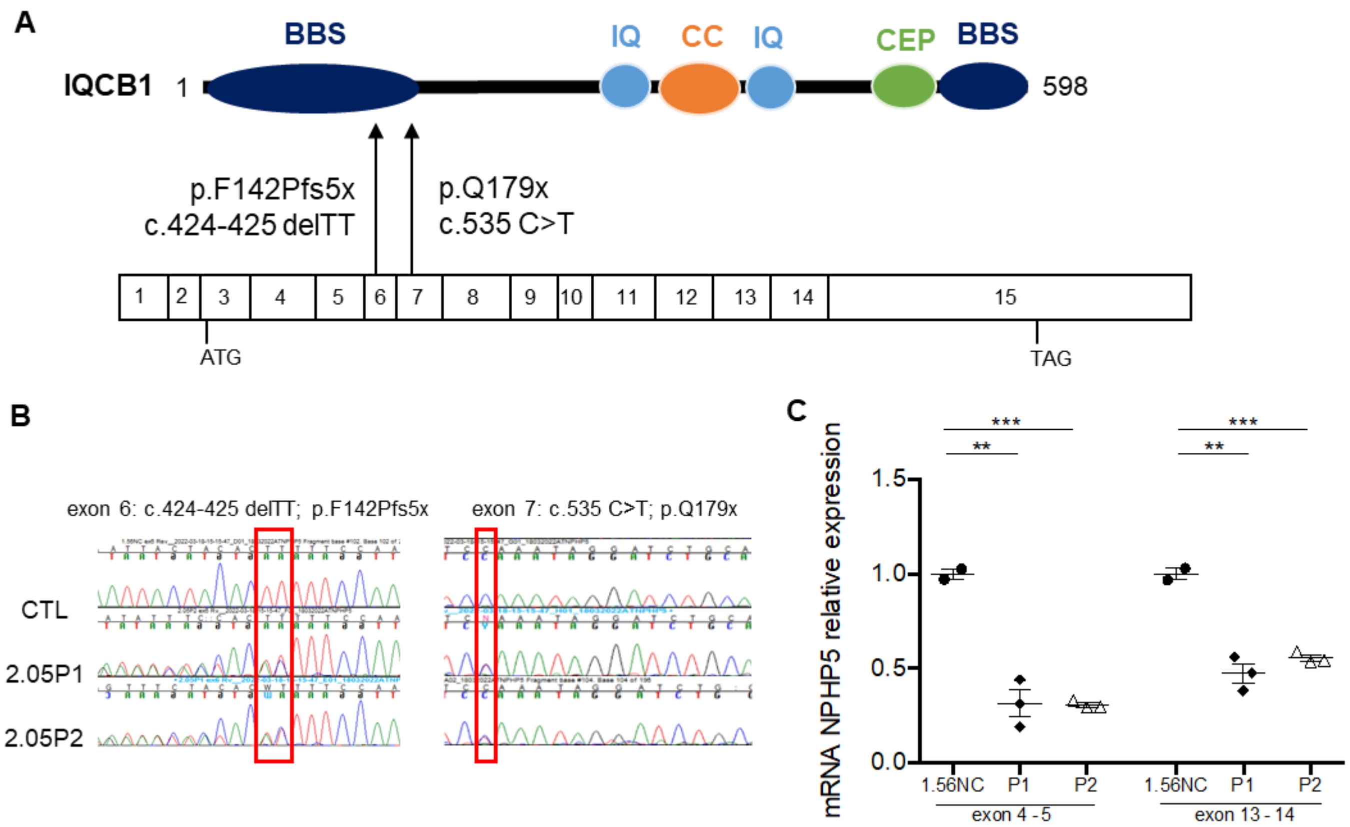
A
IQCB1 1
BBS
IQ
CC
IQ
CEP
BBS
598
p.F142Pfs5x
c.424-425 delTT
p.Q179x
c.535 C>T
1 2 3 4 5 6 7 8 9 10 11 12 13 14 15
ATG
TAG
B
exon 6: c.424-425 delTT; p.F142Pfs5x
exon 7: c.535 C>T; p.Q179x
CTL
2.05P1
2.05P2
C
mRNA NPHP5 relative expression
1.5
1.0
0.5
0.0
***
**
1.56NC
P1
P2
exon 4 - 5
exon 13 - 14

**Figure S7:**

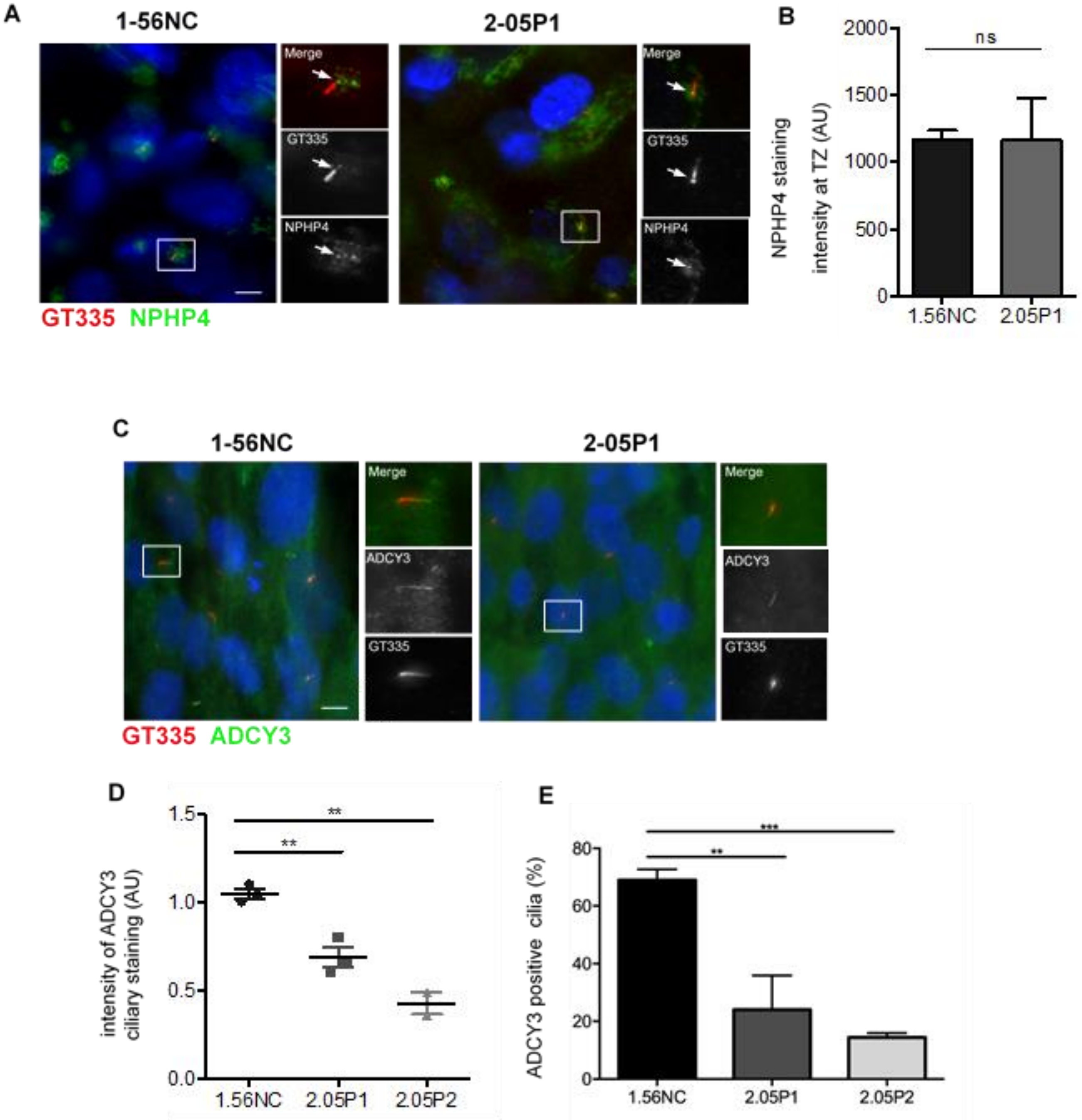
A
1-56NC
2-05P1
Merge
GT335
NPHP4
GT335 NPHP4
B
NPHP4 staining intensity at TZ (AU)
ns
2000
1500
1000
500
0
1.56NC
2.05P1
C
1-56NC
2-05P1
Merge
ADCY3
GT335
GT335 ADCY3
D
intensity of ADCY3 ciliary staining (AU)
**
**
1.5
1.0
0.5
0.0
1.56NC
2.05P1
2.05P2
E
ADCY3 positive cilia (%)
***
**
80
60
40
20
0
1.56NC
2.05P1
2.05P2

**Figure S8:**

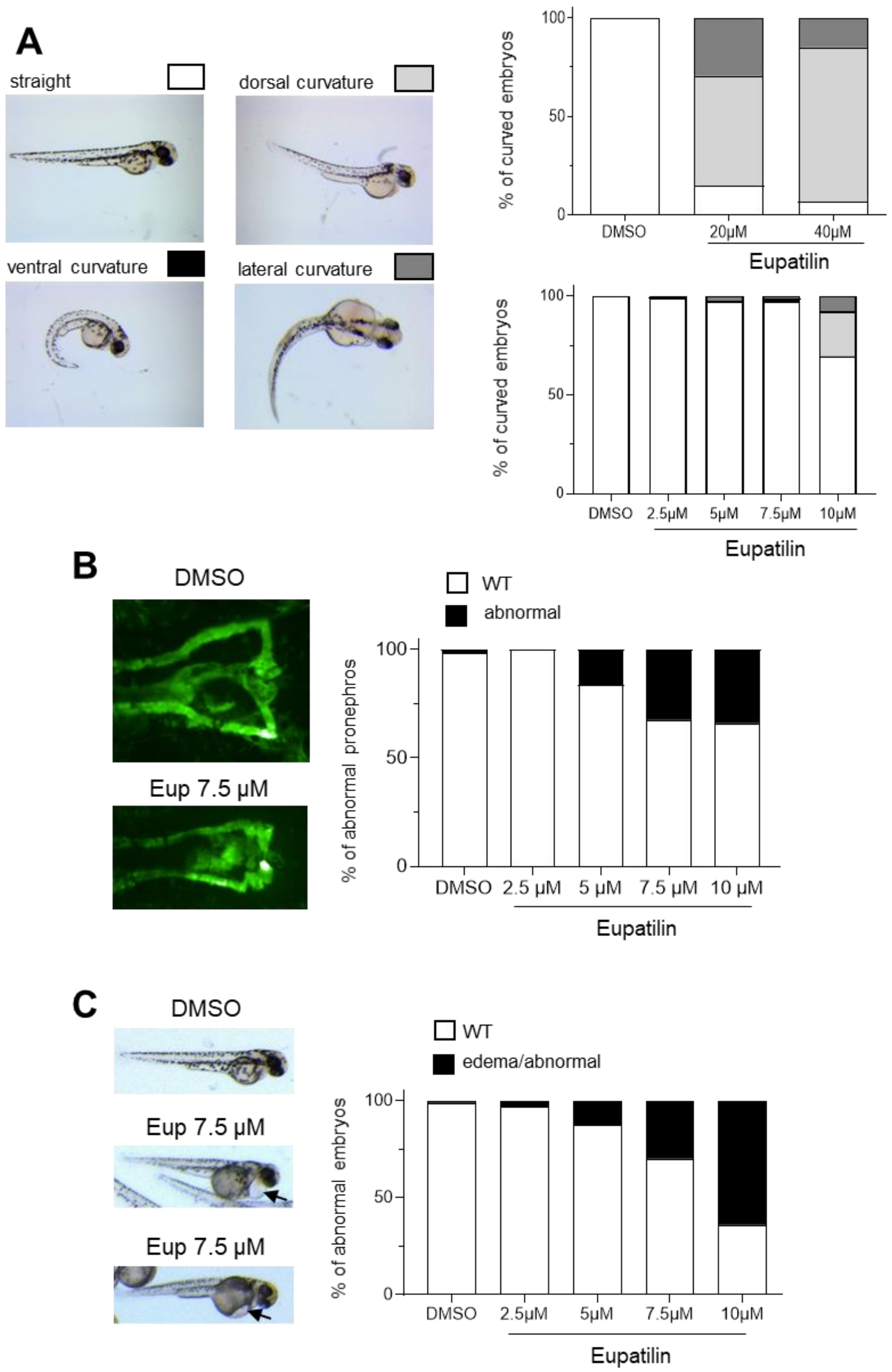

A
straight
dorsal curvature
ventral curvature
lateral curvature
% of curved embryos
100
50
0
DMSO
20µM
40µM
Eupatilin
% of curved embryos
100
50
0
DMSO
2.5µM
5µM
7.5µM
10µM
Eupatilin
B
DMSO
Eup 7.5 µM
WT
abnormal
% of abnormal pronephros
100
50
0
DMSO
2.5 µM
5 µM
7.5 µM
10 µM
Eupatilin
C
DMSO
Eup 7.5 µM
Eup 7.5 µM
WT
edema/abnormal
% of abnormal embryos
100
50
0
DMSO
2.5µM
5µM
7.5µM
10µM
Eupatilin

**Supplementary Table 1:**

| Gene Name | Sequence |
|---|---|
| CDKN2B-F | CACCCCCACCCACCTAATTC |
| CDKN2B-R | TGAGTGTCGAGGGCCAGATA |
| WDR35-F | CAACTCTAGCCGTCTTGCTATC |
| WDR35-R | TCTTTGGCCCACTTCATATCC |
| KIF3A-F | GTGGACCTTTCTCACGTGTATC |
| KIF3A-R | CGCTTTCTTCTCCCTGTCTTT |
| KIF3B-F | GGAGAACCAGCAGATGATGAA |
| KIF3B-R | GTCCAGCTCTAACAGCACAA |
| IFT88-F | TGGCAGAAGACCTCCAATAAC |
| IFT88-R | CTGCTGTCATGGGTCTAGTAAC |
| WDR19-F | ACACATTGTACCCATCCTGAC |
| WDR19-R | GACCATTCCCTCGATCTTCTTT |
| SERINC1-F | CAGCATCCGTACTTCAAACAATAG |
| SERINC1-R | TGACACCATCCCTTTCATTATCT |
| LARP1-F | CGCCAAAGAAGGCTACAGATA |
| LARP1-R | CAGAACTTCTCCAGCCCATAC |
| CD63-F | GTTTGCCATCTTTCTGTCTCTTATC |
| CD63-R | ATCGAAGCAGTGTGGTTGT |
| ILF2-F | GCAGGACATGGTCTGCTATAC |
| ILF2-R | CTTCTTCTCTGGTGGCTTCTC |
| KIF22-F | AAGAACTGGAGGCCAAGATG |
| KIF22-R | GCTGCCTGCTCCTGAATTA |
| NUP188-F | GTGGATGTCATTGCTTCTTGTG |
| NUP188-R | TTCATCCCTTCCGCACTAATC |
| TCOF1-F | CCAAGGCAGAGACAGAGAAAG |
| TCOF1-R | CAGTTTCTGAGACCAACGTAGT |
| TP53-F | GGAAGAGAATCTCCGCAAGAA |
| TP53-R | CACGGATCTGAAGGGTGAAATA |
| MYBL2-F | CACTACCAGGACACAGATTCAG |
| MYBL2-R | GTCAGTGCGGTTAGGGAAG |
| RB1-F | CACAACCCAGCAGTTCGATA |
| RB1-R | CAACATGGGAGGTGAGAGTTT |
| RBP4-F | GACAGCTACTCCTTCGTGTTT |
| RBP4-R | GTAACCGTTGTGGACGATCA |
| MAOA-F | CAGGAACGGAAGTTTGTAGGT |
| MAOA-R | TTCATGGTTCAGCGTCTCTATG |
| IGF2-F | CGTGCTTCCGGACAACTT |
| IGF2-R | CGTTTGGCCTCCCTGAAC |
| GPX3-F | TCCTTCCTACCCTCAAGTATGT |
| GPX3-R | AGAAGAGGCGGTCAGATGTA |

| | |
|---|---|
| TFPI2-F | GGCCCTACTTCTCCGTTACTA |
| TFPI2-R | TTTCTATCCTCCAGCAAGCATC |
| GALNT15-F | TCTGGAAAGCTCCACAACAC |
| GALNT15-R | GCCAAAGTGAATCTCCTTCCT |
| ATOH8-F | TGACTACAGTGCCGACCACA |
| ATOH8-R | TCACTCCTTGCGCTTCTTGG |
| PDK4-F | GGTGGTGTTCCCCTGAGAAT |
| PDK4-R | GGCAAGCCGTAACCAAAACC |
| LAMC2-F | CGCAGCTCTGCAGAATACAG |
| LAMC2-R | AGACCCATTTCGTTGGACAG |
| CAV1-F | CCAAGGAGATCGACCTGGTCAA |
| CAV1-R | GCCGTCAAAACTGTGTGTCCCT |
| PKIA-F | TGATATCCTGGTTTCCTCTGCA |
| PKIA-R | GGCTTCCCCACTTTGTTCTG |
| TUBB-F | GAAGCCACAGGTGGCAAATA |
| TUBB-R | CCCAGACTGACCAAATACAAAGT |
| MYLK-F | CAACCTCACTGTCGTGGATAAG |
| MYLK-R | TCCCAGATCTCGATGCTGTA |
| ACTN1-F | TGGGTTACAACATGGGAGAAG |
| ACTN1-R | CATGACTTGGTCTGCTGTATCT |
| LAMB1-F | ACTGTTTCCAGGGAGTGTATG |
| LAMB1-R | CCAAGCACCTTTCACAGTTATG |
| CDK4-F | GAAGTTCTTCTGCAGTCCACATA |
| CDK4-R | CAGCCCAATCAGGTCAAAGA |
| CDK2-F | GAGTTGTGTACAAAGCCAGAAAC |
| CDK2-R | ACATCCAGCAGCTTGACAATA |
| PCNA-F | AGGAGGAAGCTGTTACCATAGA |
| PCNA-R | AGTGTCCCATATCCGCAATTT |
| NPHP5-exon4-5-F | TGCCTCTTGGTCCTCAGTCA |
| NPHP5-exon4-5-R | GCCCACACAGCAATGGCTTA |
| NPHP5-exon13-14-F | TCAGAAGACATTTGGGCTCTCC |
| NPHP5-exon13-14-R | TCGTTCTTGAGCTTGGGCAT |
